# Separable cognitive and motor decline tracked across adult life-span for goal-directed navigation

**DOI:** 10.1101/2024.12.03.626546

**Authors:** Gian Luca Lancia, Marco D’Alessandro, Mattia Eluchans, Miguel Ibáñez-Berganza, Hugo J. Spiers, Giovanni Pezzulo

## Abstract

Spatial navigation is a fundamental cognitive skill, yet our understanding of the strategies people employ in real-world navigation remains incomplete. This study investigates navigation strategies using data from the large-scale Sea Hero Quest (SHQ) game, in which participants face navigational problems consisting in controlling a boat to reach a series of goal locations, within maps having different shapes. By using a combination of behavioral and computational modeling approaches, we report three key findings: First, participants predominantly use goal-directed strategies compared to visibility-based strategies. Second, participants show signatures of sequential planning, both during memorization – as they spend more time looking at difficult levels – and during navigation – as they show primacy effects, suggesting that they remember better the first navigational goal. Third, our results show a significant age-related decline in navigation performance, which depends both on reduced motor skill – as indexed by poorer control of the boat – and decreased cognitive strategy – as indexed by a loss of goal-directedness during spatial navigation. Unexpectedly, the loss of goal-directedness with age is not associated with a shift to visibility-based strategy or to an incorrect sequencing of goals. Taken together, our findings offer a nuanced understanding of spatial navigation strategies in naturalistic conditions and their decline with age. Our computational approach permits distinguishing motoric and cognitive variables, showing that both decline with age. These results align with and extent previous research on spatial memory decline with age, including potential links to neurodegenerative diseases like Alzheimer’s. The computational models developed in this study provide a robust framework for analyzing navigation strategies, applicable to other virtual environments and video games.

## 1 Introduction

Spatial navigation is a fundamental and ubiquitous skill for humans and other animals (Epstein et al., 2017). Human navigation and wayfinding strategies have been extensively studied but the large majority of studies focused on relatively small scale settings (with few notable exceptions (Bongiorno et al., 2021)). Hence, we still have an incomplete view of how people make navigation decisions in real life conditions.

In recent years, there has been a growing interest in the use of video games to study human spatial navigation (Allen et al., 2023). In game-like settings, participants typically explore virtual environments by controlling an avatar, in first or third person perspectives. Using game-like settings that mimic some real-world properties — such as spatial organization — might help preserve some ecological validity and engagement, while at the same time allowing the experiments to be conducted remotely. In this sense, a very successful example is the Sea Hero Quest project (SHQ): a spatial navigation video game that was played by more than 6 million participants (Spiers et al., 2023; Coutrot et al., 2019, 2018a; West et al., 2023). SHQ can be considered a task in which cartographic map information is presented for memorization and must be used to guide first-person navigation in ocean/river based environments. Past research comparing SHQ to a city version of the task involving car driving has found very comparable effect of background demographics (Coutrot et al., 2022). In SHQ, participants face a sequence of navigation problems (or levels). For each level, they first memorize the problem map (presented in allocentric perspective) and then they control a simulated boat (in third-person perspective) to solve the problem, which consist in reaching three or four navigational goals in the correct sequence, as soon as possible (Figure 1).

**Figure 1:**
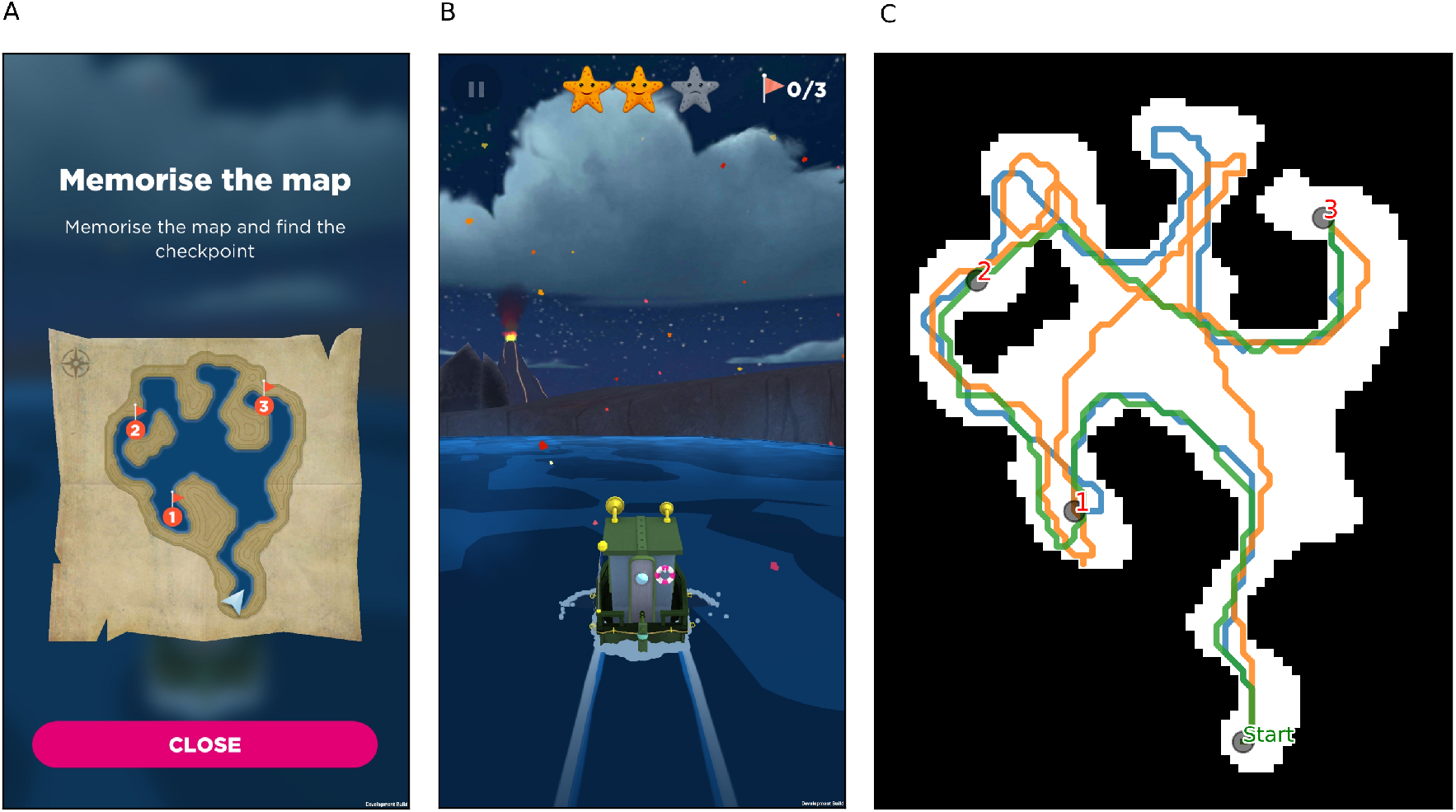
The Sea Hero Quest game. A) At the beginning of the level, players can see and memorise the map in an allocentric view. B) An example of the third-person view navigation. Players can change the direction of the boat by tapping at its left or right. The stars at the top of the screen give a visual cue of how much time remains. C) The trajectories of three players (blue, green and orange) in level 46 are shown as examples. Note that environmental landmarks such as the volcano in (B) are not shown in the map in the allocentric view

The analysis of SHQ data revealed that there are large differences between navigation abilities across age groups, with elder participants showing worse navigation performance compared to younger participants (Spiers et al., 2023), as well as differences with gender, education level and environment where they live in (Coutrot et al., 2018b). These results transfer to the real world, as the performance in SHQ was found to be correlated with the performance in real world navigation tasks (Coutrot et al., 2019). Additionally, the analysis of SHQ data revealed that the game is able to detect the presence of a genetic predisposition to Alzheimer’s disease (Lim et al., 2023; Coughlan et al., 2019), and thanks to the large number of participants, it was possible to show that the handedness of the participants is not correlated with their navigation performance (Fernandez-Velasco et al., 2023).

However, these studies considered relatively simple behavioral measures of performance, such as the percentage of completed problems and the time taken to complete them. While important, these synoptic analyses cannot reveal what navigation strategies participants adopt to solve SHQ problems and whether the performance decrease found across age groups depend on poorer cognitive processing (e.g., a lack of goal-directedness, possibly reflecting poor spatial memory) or poorer motor control (e.g., the inability to control the boat accurately). These questions are particularly pressing in the context of SHQ, since the game was designed with the goal to provide ecologically valid measures of spatial memory and of its decline with age.

Computational modeling of behavior is an appealing way to complement behavioral measures and to probe the mechanisms underlying human navigation abilities. Existing computational models of navigation range from simple rule-based systems to complex neural network architectures that can mimic human cognitive processes. For instance, models of cognitive mapping and path integration simulate how individuals navigate by encoding spatial information and updating their position relative to known landmarks (Tolman, 1948; Gallistel, 1990). Other models addressing wayfinding incorporate elements of decisionmaking and strategy selection, allowing researchers to predict how different environmental cues and cognitive strategies influence navigation behavior (Burgess, 2008; McNamara et al., 2003). These models are invaluable for dissecting the cognitive and neural bases of navigation, offering insights into how different factors such as age, experience, and environmental differences affect navigation performance. Moreover, “lesioning” the models permits examining the effects of individual cognitive processes on navigation performance (Zhi-Xuan et al., 2020; Lancia et al., 2023; Jacob et al., 2023).

One influential class of models describes planning—whether spatial or not—as a form of mental simulation used to evaluate possible future trajectories (Daw and Dayan, 2014; Hunt et al., 2021; Gelly and Silver, 2011). A key limitation of this approach is that the number of trajectories to simulate increases rapidly, even in simple contexts, necessitating heuristics or approximate strategies to manage the computational load (Kolling et al., 2018; Huys et al., 2012, 2015). An alternative perspective suggests that individuals do not simulate all possible futures from scratch. Instead, they rely on a *default policy*—a habitual or reference behavior in the given context—from which planning diverges as needed to meet task demands, as formalized in (Todorov, 2006; Piray and Daw, 2020; Rubin et al., 2012a) and empirically tested in (Lancia et al., 2023). However, identifying which *default policy* is engaged in a particular task remains challenging. Inspired by this second class of computational models for spatial navigation we developed a set of computational models to solve the navigation tasks of SHQ arbitrating between a goal-directed strategy aimed at reaching the next (sub)goal as efficiently as possible, and a default strategy favoring the direction of maximal environment visibility— both of which have been consistently implicated in spatial navigation (Mcelhinney et al., 2022; Turner and Penn, 2002; Mattar and Lengyel, 2022; Eluchans et al., 2023; Emo, 2014). We used these computational models, alongside model-free analyses, to assess participants’ goal-directedness and its decline with age in SHQ.

Specifically, in this study we addressed three main questions. First, we asked whether participants privileged goal-directed versus visibility-based strategies during navigation and how the balance of the two strategies is modulated by map characteristics, such as goal visibility. Second, given that SHQ involves planning and remembering sequences of navigation goals, we asked whether we could identify signatures of sequential planning both when participants memorize the problem maps and when they later navigate them. Third, and most importantly, we asked whether we could observe an age-dependent decline in navigation skill and whether our computational models could help assess whether it could be due to a decline of motor skill (e.g., decreased ability to control the boat), cognitive strategy (e.g., decrease of goal-directedness, shift from goal-directed to visibility-based strategies), or a combination of both factors.

## 2 Methods

### 2.1 Experimental setup: The Sea Hero Quest game

In Sea Hero Quest, participants of all ages navigate in a virtual environment by controlling a boat in third person view with a smartphone or tablet (see Figure 1B). By tapping left or right on the screen, they can change the direction of the boat, and by swiping in the forward direction they can increase the speed. The goal of the game is to reach a series of checkpoints as fast as possible, while collecting as many golden stars as the limited time allows, up to a maximum of 3 or 4 stars depending on the level. An allocentric and complete view (top view) of the map and of the goals positions in it was shown before starting the navigation for as long as the player wanted (see Figure 1A).

The dataset used in this study was collected from the Sea Hero Quest game, a mobile game developed by Glitchers in partnership with Deutsche Telekom, University College London, and the University of East Anglia. Designed to gather data on human spatial navigation, the game specifically aimed to study the navigational behavior of individuals with a genetic predisposition to Alzheimer’s disease. Since its release in 2016, Sea Hero Quest has been engaged by millions of users globally. The dataset employed for this analysis was extracted from the game’s servers, encompassing detailed records of participants’ navigational behavior, including their trajectories, the duration to achieve each goal, and their age.

The game’s mechanics involve navigating a boat through a series of virtual environments, with the objective of reaching a sequence of checkpoints (goals) as quickly as possible. Participants control the boat’s direction and speed using a smartphone or tablet, and are presented with a map of the environment at the start of each level. The dataset used in this study comprises navigational data from *n* = 18456 participants, who completed all 17 levels that we used. Thus, the dataset includes a total of 313752 unique trajectories, recorded at a resolution of 1 Hz, with x and y coordinates and the orientation of the boat. These 17 problems were selected because, among the various tasks included in the full SHQ game, some did not involve sequential navigation from goal to goal—for example, tasks where participants had to reach a final location and then indicate the direction of the starting point. These non-sequential tasks were excluded. From the remaining problems, we also filtered out any cases where the map shown at the beginning of the trial included occlusions. Among the valid problems, we chose the first 17 encountered in the game because they had the largest sample sizes. While navigation is in a continuous space, the trajectories are recorded into a discretized grid, with each cell representing a state, and the orientation angles are divided into 16 possible angles, which we further reduced to 8. Additionally, we interpolated the trajectories to avoid movements between non adjacent cells. We used this dataset in all the analyses which don’t explicitly specify the use of age as a variable.

Of the participants, *n* = 14612 completed the survey, providing their age, gender and other information such as handedness and country of origin, and are all aged between 16 and 99 years. As in previous studies on this dataset (Spiers et al., 2023), we excluded participants younger than 19 years old, as well as those older than 70 years old, to avoid the influence of extreme age groups. As shown in (Spiers et al., 2023), instead of a decline in performance, after 70 years old it was found an improvement. This counterintuitive result is likely due to a selection bias in that population, thus, we limited the age of the participants used in this study between 19 and 70 (*n* = 5896). The cohorts used in this study are thus 19-35 (*n* = 1766), 36-55 (*n* = 2719) and 56-70 (*n* = 1411) years old, which provide an approximately balanced sample size among the three groups. We also excluded participants who reported being 18 years old, as this was the default age presented by the application and could have been accepted by users without being actively selected. We used this dataset in all the analyses which explicitly specify the use of age as a variable.

Each problem (or level) is characterized by a unique grid layout, with varying numbers of goals and obstacles, and participants’ trajectories are recorded as they navigate through these environments. Levels used in this study are shown in Figure 2. Levels 1 and 2 are the only two levels with a single goal, and are meant to familiarize with the game and are thus excluded from our analysis; levels 26, 27, 32, 42 and 43 are the only ones that we included with 4 goals. The other levels have 3 goals.

**Figure 2:**
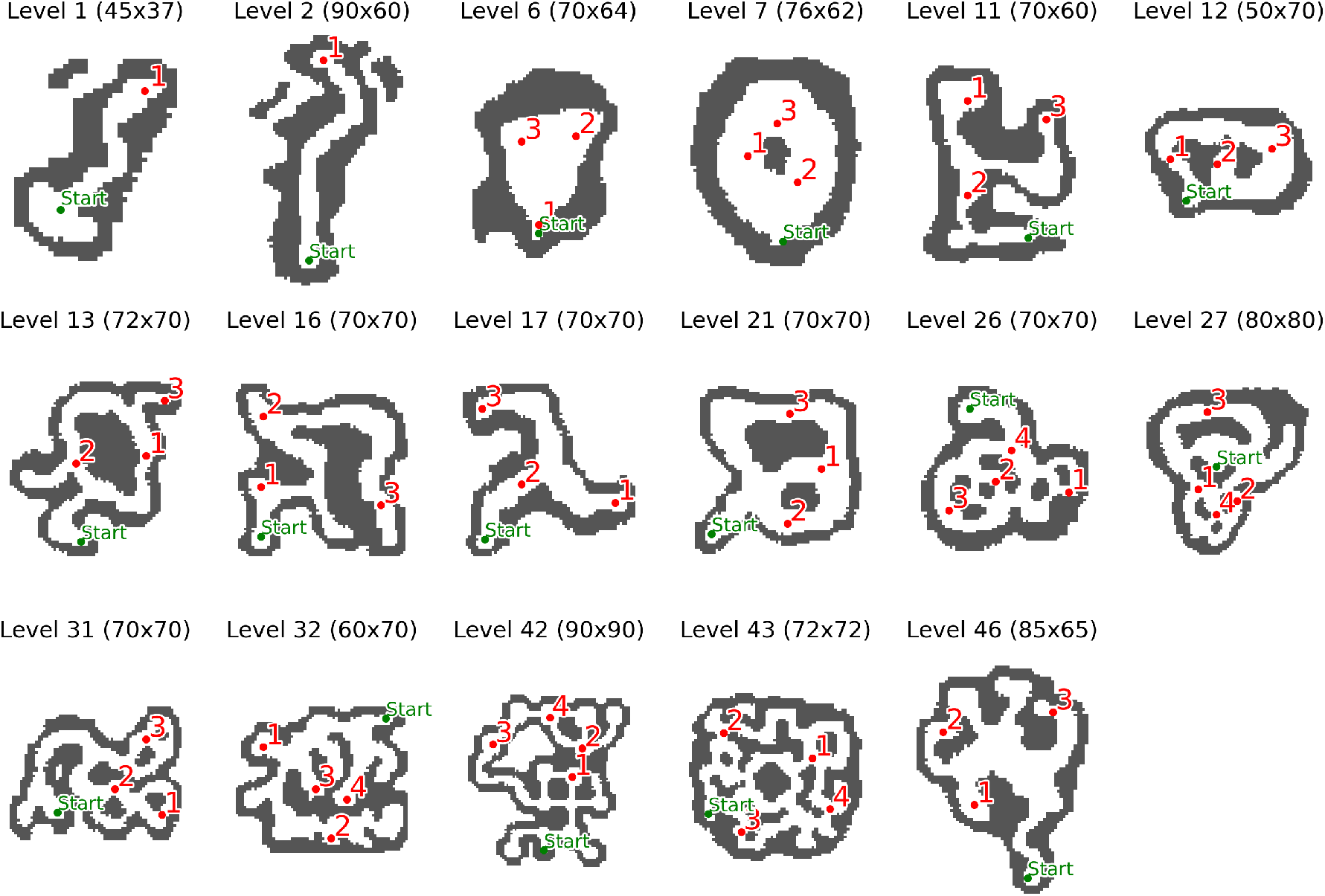
The 17 levels (or problems) of Sea Hero Quest included in our analysis. The title shows the level number and size of the grid (height and width). In each map, the Start is represented by a green circle, the Goals by red circles, the walls are dark grey. Level 1 and 2 are the only two levels with a single goal, and are meant to familiarize with the game; levels 26, 27, 32, 42 and 43 are the only ones that we included with 4 goals. The other levels have 3 goals.

### 2.2 Computational modeling of navigation

Navigation is a complex process that involves a number of cognitive and perceptual processes, however we aimed to develop simple models that could try to capture some of its main aspects in the game. We developed two types of models based on the premise that participants navigate by integrating their perception of the environment with their objectives.

The first type, termed Visibility-based model, draws inspiration from previous research on the exosomatic visual agent system (EVAS) (Mcelhinney et al., 2022; Turner and Penn, 2002), positing that navigation is primarily influenced by the perception of environmental affordances. Affordances, in this context, refer to the action possibilities offered by the environment, relative to the agent’s abilities. In natural settings, individuals navigate by avoiding obstacles and seeking vantage points to enhance visibility, often choosing paths that maximize environmental visibility (Turner and Penn, 2002). This approach to navigation does not account for explicit goals but rather navigates by optimizing the visibility of the surrounding environment. Thus, the more participants follow visibility affordances, the higher the likelihood of the Visibility-based model.

The second model type, referred to as the Goal-Directed model, characterizes agents following the shortest path to the correct goal. Thus, the better a participant approximates the shortest path to the correct goal, the higher the likelihood of the Goal-Directed model. While following the shortest path might seem a strong criterion, it is important to remind that in SHQ, participants are shown goal locations in the map at the beginning of each level, potentially allowing them to plan the shortest route to the goals. Yet, errors in memory recollection may cause the selection of suboptimal paths that differ from the shortest path to the correct goal (or go towards an incorrect goal) – which imply a lower likelihood of the Goal-Directed model.

We used these models to simulate the navigation of participants in the game. Figure 3 illustrates the behavior of the model, by qualitatively comparing them with experimental data (i.e., the actual navigation of participants) in an example level (Level 46). Looking at the experimental occupancy (Figure 3A) we can see that players tend to occupy the states included in the shortest routes from Start to Goal 1 (blue), then from Goal 1 to Goal 2 (green) and finally from Goal 2 to Goal 3 (orange). This is consistent with the idea that participants navigate by following a goal-directed strategy to reach their current goal. However, there are areas of the map with a high occupancy that are not part of the goal-directed strategy, for example the upper alcove in between Goal 2 and 3. This suggests that participants navigate by integrating both goal-directedness and some other strategy, possibly related to the visibility of the environment. In this particular case, a possible explanation of this behaviour is that participants move in the direction of the goal and when presented with a wide open area on their left, they deviate towards that direction. This suboptimal path cannot be explained by the simulated goal-directed strategy (Figure 3B, orange). On the other hand, the Visibility-based model is not able to capture the trajectories of participants when taken alone (Figure 3C): the Visibility-based model lacks the explicit objective of going towards the goals, and thus it is not able to solve the task within time limits. However, as can be seen by the occupancy map, this simple strategy allows the agent to explore the map more thoroughly in absence of explicit goals, and could reflect the behavior adopted by participants when they are not actively pursuing a goal, such as for example when they get lost.

**Figure 3:**
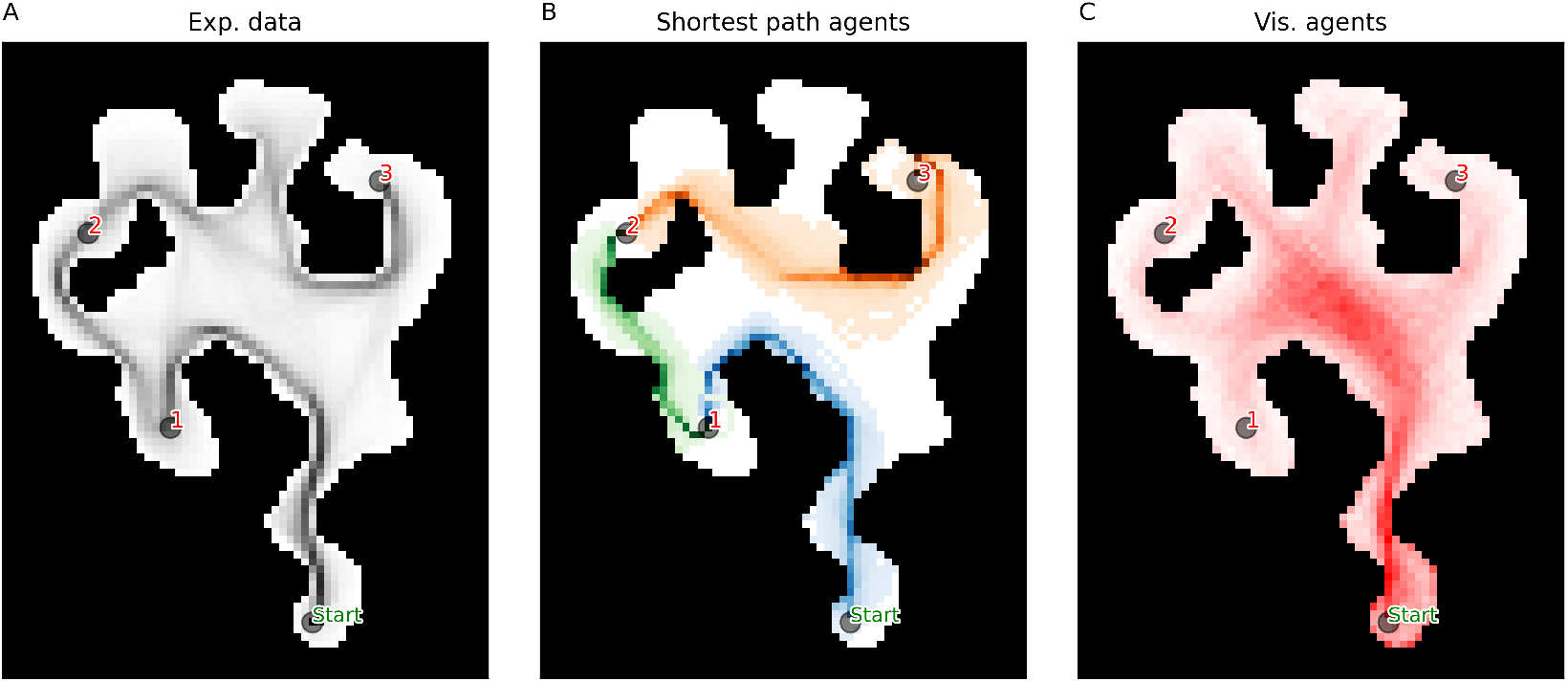
Comparison between the occupancy of participants and those of Goal-Directed and Visibility-Based models. Occupancy maps of: A) n=1000 participants randomly chosen, a darker hue of black means a higher occupancy. B) *n* = 200 Goal-Directed models, with the inverse temperature parameter *β*, that governs the randomness of the model policy, uniformly distributed between 1 and 5; for each run the model started with Goal 1 as goal, after reaching it it shift its objective to Goal 2 and then to Goal 3. C) *n* = 200 Visibility-based models with *β* uniformly distributed between 1 and 5, for a maximum of *T* = 500 steps.

We provide an illustrated example of the qualitative comparison between the navigational behavior of individual participants with model predictions in Figure 4. Figure 4A shows the complete trajectory of a participant navigating Level 46, tracing their movement from the Start point to Goal 3. The color coding at each trajectory point indicates the model with the highest likelihood of predicting the participant’s action at that step (i.e., each state-action pair), with the red color representing the Visibility-based model and the blue, green and yellow colors representing the Goal-Directed models for Goals 1, 2 and 3, respectively. Figure 4B-E shows the same results but split by segment (from Start to Goal 1, from Goal 1 to Goal 2, and from Goal 2 to Goal 3). The analysis shows that for each segment, the “correct” Goal-Directed model has the highest likelihood, indicating that participant’s navigation is largely goal-directed. Note that in the initial part of the first segment, the first two Goal-Directed models and the Visibility-based model have a high likelihood, indicating that not the all the segments of all the maps allow full distinguishability of the models. The highest likelihood of the Visibility-based model in the very first part of the segment is likely due to map characteristics, with only one path being available.

**Figure 4:**
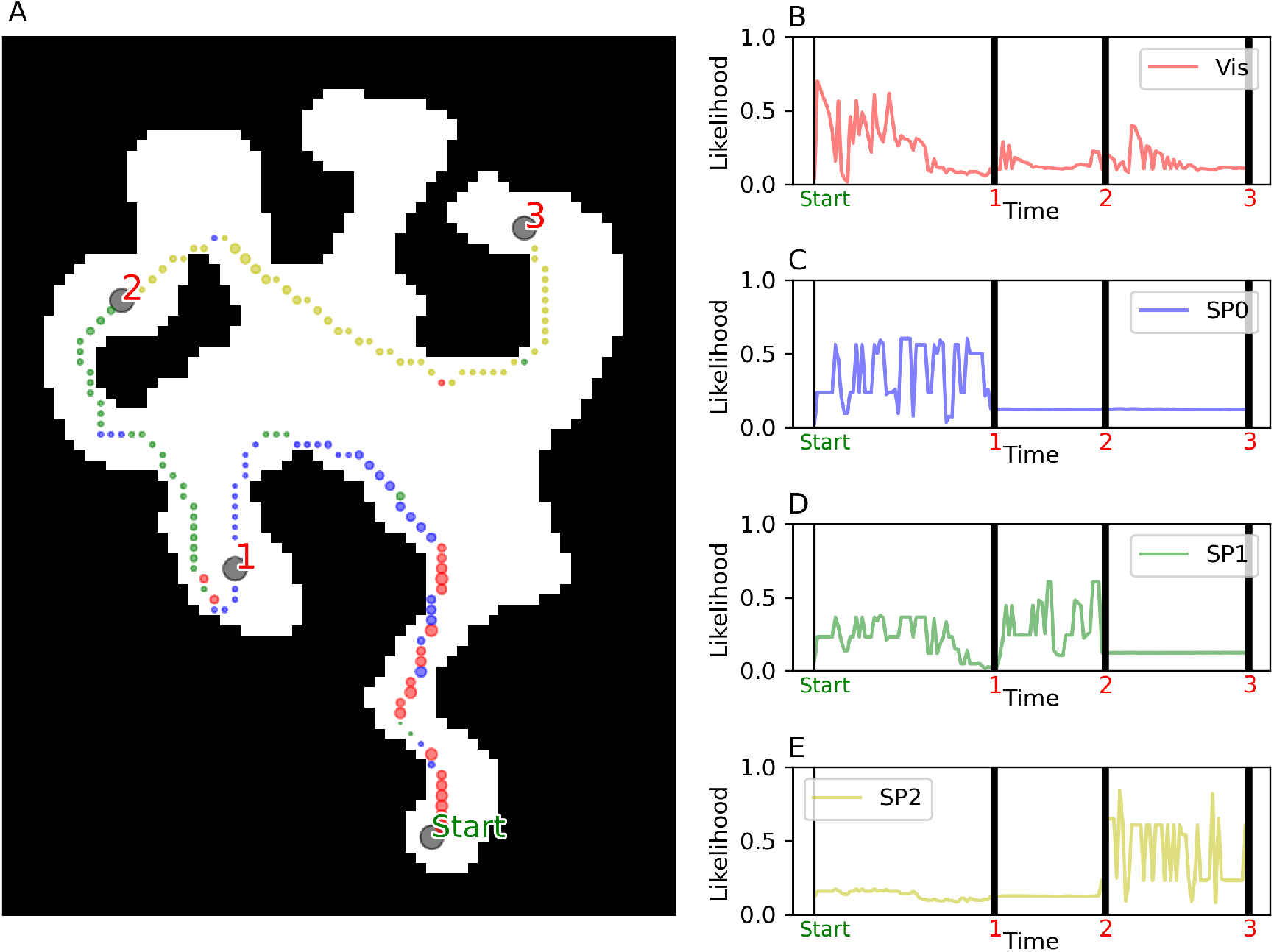
An example illustrating our model-based analysis. A) Trajectory of a participant navigating in Level 46, from Start to Goal 3. The color of each trajectory point displays which model has the highest likelihood at that particular step (i.e., state-action pair), with the red color representing the Visibility-based model and the blue, green and yellow colors representing the Goal-Directed models for Goals 1, 2 and 3, respectively. B-E) The four plots display at each time step the likelihood of the action taken by the participant given the four models: Visibility (Vis), Goal-Directed model for Goal 1, Goal 2 and Goal 3 (respectively GD 1, 2 and 3). Black vertical bars show the time step when a Goal is reached.

### 2.3 Formal specification of the computational models

In this section we describe the models used to simulate the navigation of participants in the game, and how we used these models to evaluate the data and simulate new data. We used two types of models, the first one is based on the idea that participants navigate by integrating their perception of the environment, that is, they tend to avoid obstacles and follow the directions that offer greater visibility- or, in other words, those that lead towards wider open areas; the second one is based on the idea that participants navigate by identifying and following the most efficient path to a predetermined goal, in other words they use some kind of internal map of the environment to plan their route.

To formalize these models, the Markov Decision Process (MDP) framework was employed (Puterman, 1994; Bellman, 1958). This mathematical framework delineates the process of decision-making, where a decision-maker engages in a series of choices. Each choice influences the state of the environment and the ensuing rewards.

In the context of Sea Hero Quest, the environment is a grid, with each cell representing a state *s*, and the agent can move from one state to another by taking an action, following the rule of a deterministic transition matrix *P* (*s*^*′*^|*s, a*), which gives the probability of reaching state *s*^*′*^ from state *s* by taking action *a*. Only transitions towards adjacent states are allowed. The agent takes actions following a policy *π*(*a*|*s*), which gives the probability of taking action *a* in state *s*. Both the goal-directed and the affordance-based models are defined by a policy, and they differ for the way the policy is obtained.

#### 2.3.1 Visibility-based model

The visibility-based (Vis) model is defined by a policy that gives the probability of taking an action in a state given the visibility of the environment from that state. All the states in direct line of sight (e.g., not occluded by walls) are divided into 8 bins, each spanning an angle of 45° (see Figure 5A). In this way, we obtain a histogram of the number of states in each bin *n*(*s, a*) (with *s* representing the states the agent is in, and *a* the bin, see Figure 5B), and the policy is defined by the probability of taking an action in a bin given the number of states in that bin. The policy is defined by a softmax function, with the probability of taking an action in a bin given by the number of states in that bin, and a parameter *β* that controls the randomness of the policy (see Figure 5C). The policy is defined as:

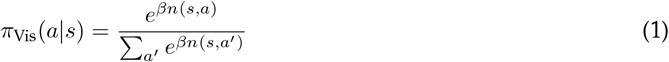

**Figure 5:**
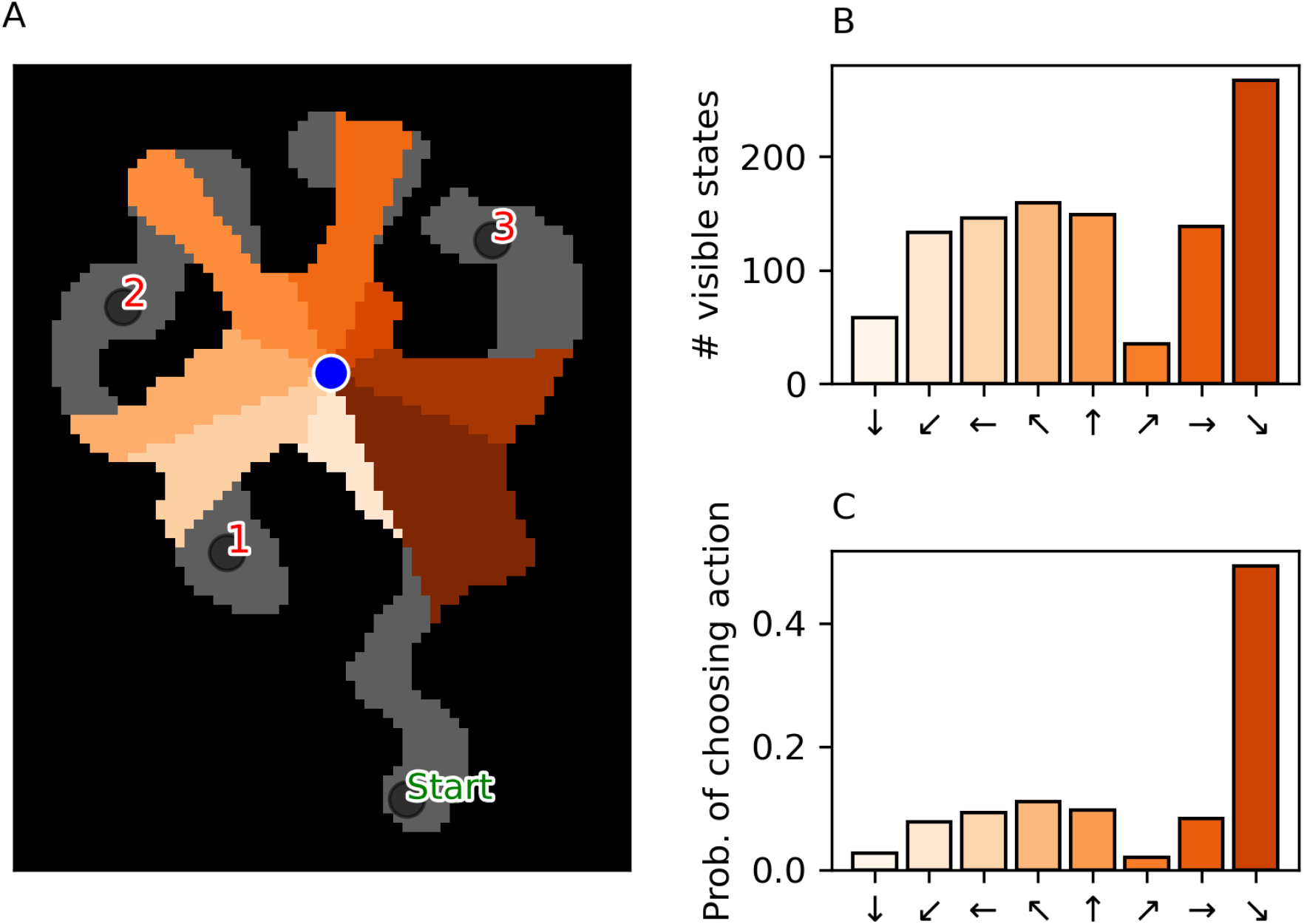
An example illustrating the Visibility-based model. A) Area of the map visible from the point of view of a simulated player (blue circle) in a given position, of level 46 from the center of the map. The visible area is subdivided in 8 bins, each spanning an angle of 45°. Grey areas are not visible. B) Histogram showing the number of states in each cone. C) An example of softmax function applied to the number of states visible in a bin to obtain the probability of choosing the related action. See main text for explanation.

The denominator of the softmax ensures that the probabilities sum to 1. The parameter *β* controls the randomness of the policy, with lower values leading to more random policies. The policy is defined in this way to reflect the idea that the agent is more likely to take an action in a bin with more states.

#### 2.3.2 Goal-Directed model(s)

A Goal-Directed (GD) model is defined by a policy *π*(*a*|*s*) that returns the probability of taking an action *a* from state *s* to state *s*^*′*^, given a distance *d*(*s*^*′*^, *g*_*i*_) between state *s*^*′*^ and a goal *g*_*i*_. Given that SHQ problems involves 3 or 4 goals, for each problem there are either 3 or 4 alternative Goal-Directed models, one for each segment (see Figure 4).

Each model favours actions that minimize the distance *d* to its goal, and is defined as:

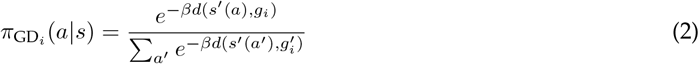

where *i* refers to the goal *g*_*i*_ the agent is currently pursuing, and *β* is a parameter that controls the randomness of the policy.

#### 2.3.3 Model fitting and comparison

For each model, we used the policy to calculate the likelihood of the actions taken by participants in each state, and then computed the log-likelihood of the trajectories:

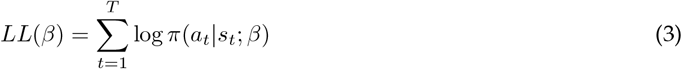

where *t* is the time step, *T* is the total number of time steps, *a*_*t*_ is the action taken by the participant at time *t*, and *s*_*t*_ is the state the participant is in at time *t*. We then used the log-likelihood to fit the only free parameter, *β*, of the policy, by maximizing the log-likelihood:

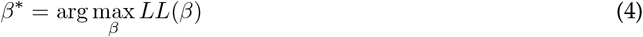

We performed model fitting using the scipy.optimize package in Python. We then used the fitted model to calculate the likelihood of the actions taken by participants in each state, and compared the likelihoods of the different models for each participants’ trajectory.

#### 2.3.4 Model simulations

In control analyses, we simulated the navigation of participants by using the policy *π*_*i*_(*a*|*s*) of each model, that is, we sampled an action from the policy *a*_*t*_ ∼ *π*_*i*_(*a*_*t*_|*s*_*t*_) to generate simulated trajectories at each time-step *t*.

We then used these simulated trajectories to calculate the likelihood *L*(*π*_*i*_) of the actions *A* = *{a*_1_, *a*_2_, …, *a*_*T*_ *}* taken by participants in each state *S* = *{s*_1_, *s*_2_, …, *s*_*T*_ *}*. The log-likelihood of *i* − *th* model’s policy *π*_*i*_ given the observed actions and states is:

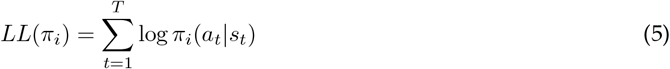

Finally, we compared the likelihoods *L*(*π*_1_), *L*(*π*_2_), …, *L*(*π*_*M*_) of the different models *π*_1_, *π*_2_, …, *π*_*M*_ for each simulated trajectory to determine which model best fits the observed data.

#### 2.3.5 Relative Goal-Directedness

We developed an index of *Relative Goal-Directedness* that quantifies the relative contribution of Goal-Directed (GD) and Visibility-based (Vis) models to explain participants’ data. We explicitly modeled the relative contributions of the two strategies by combining Equation 1 and Equation 2 into a mixture probability model that weights visibility and Goal Directed policies. The mixture policy is now defined as:

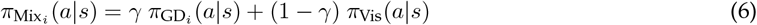

for the given goal *i*. The *γ* parameter is estimated via the same maximum likelihood procedure described above. The greater the *γ* of a participant, the higher the Relative Goal-Directedness – meaning that the participants prefers the goal-directed policy over the visibility-based policy.

## 3 Results

### 3.1 Participants consistently show goal-directed navigation

We first asked what navigation strategies participants used when solving SHQ levels. Behavioral data (e.g., navigation time) cannot easily distinguish participants’ strategies – and this motivates our model-based analysis. Previous navigation research revealed the importance of both goal-directed strategies that seek efficient paths to the current goal and visibility-based strategies that follow the direction of maximum visibility – then we designed computational models implementing both strategies. Interestingly, already the example problem showed in Figure 3 shows that the occupancy of participants (Figure 3A) is not fully accounted by a goal-directed strategy to follow the shortest path (Figure 3B), as evident from the fact that several participants visit the north branch where there is no goal. A visibility-based strategy (Figure 3C) can account for this behavior but it would predict a much greater proportion of time spent in the center of the maze, compared to what humans do. This example illustrate that both goal-directed and visibility-based strategies could be necessary to fully account for the data – prompting the question of which strategy is more prominent in SHQ.

#### 3.1.1 Goal-directed models have greater likelihood than the Visibility-based model

We asked whether a model-based analysis of participants’ navigation could reveal goal-directed navigation, over and above the Visibility-based strategy to follow the direction of maximal visibility. To address this question, we analyzed the performance of different navigational models by averaging their likelihoods across participants’ trajectories. Figure 6 illustrates these averaged likelihoods, with the models positioned along the x-axis (Visibility, Goal-Directed model to Goals 1 through 4) and the y-axis representing the models’ average likelihood based on participant actions. The models are color-coded for reference: Visibility in red, Goal-Directed model to Goal 1 in blue, Goal 2 in green, Goal 3 in yellow, and Goal 4 in azure. The two plots represent problems with 3 goals (Figure 6A) and with 4 goals (Figure 6B).

**Figure 6:**
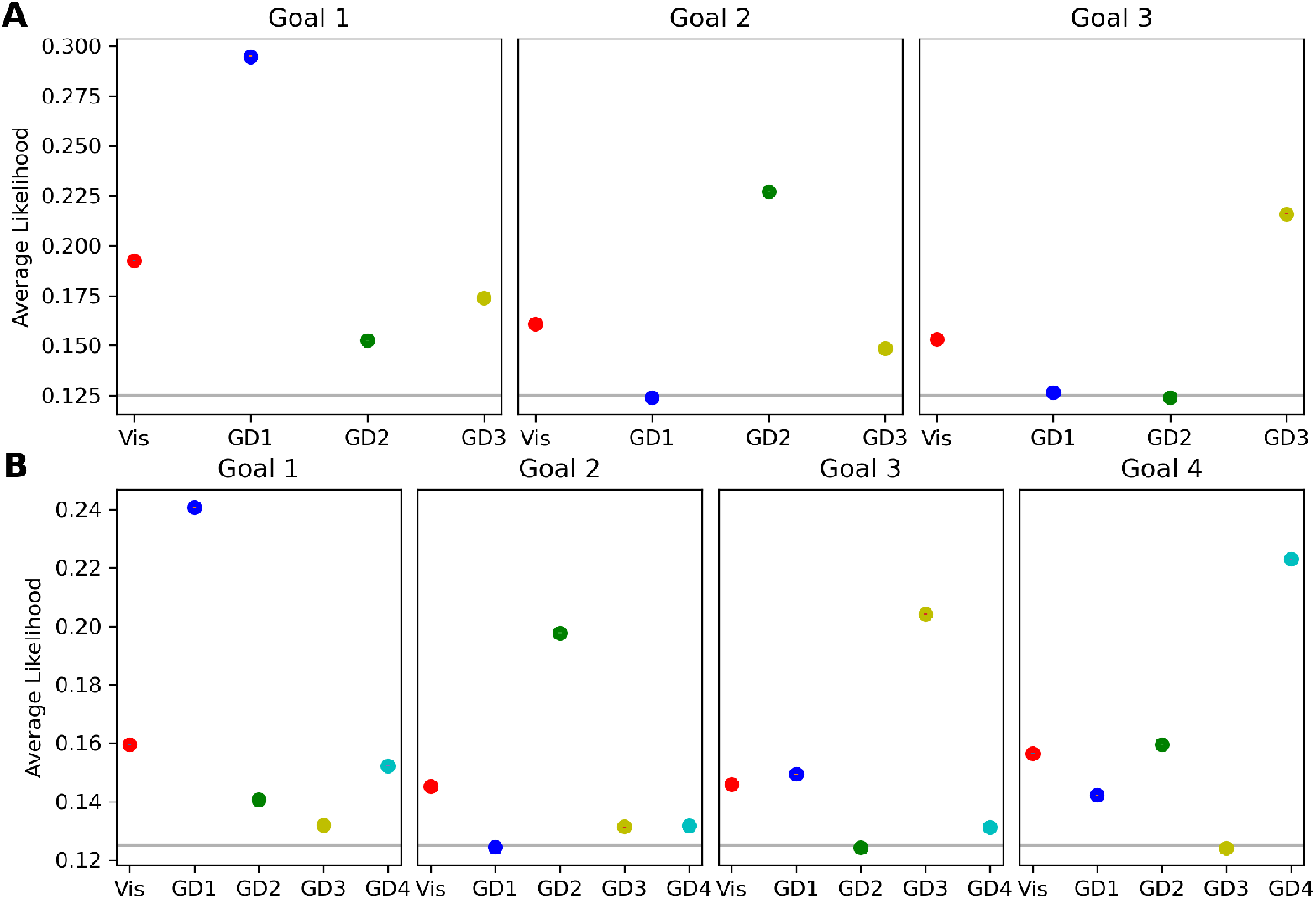
Results of the model-based analysis. Model Performance by Goal and Level. This figure presents the averaged likelihoods of the models considered in this study (Vis: Visibility-based model; GD1-GD4: Goal-Directed models 1 to 4) across participants, levels, and trajectory segments. Trajectory segments are defined by the number of goals within a level, divided into chunks of three or four, depending on the level’s goal count. A trajectory chunk ends when a goal is reached, and averages are calculated within these segments. Error bars represent standard errors across participants and levels, which are smaller than the depicted symbols. The models are color-coded: red for Visibility, blue for Goal-Directed to Goal 1, green for Goal 2, yellow for Goal 3, and azure for Goal 4. Panel A (top row) displays model likelihoods for levels with three goals, with subplots for each goal reflecting model performance as participants navigate towards each successive goal. Panel B (middle row) shows likelihoods for levels with four goals. The grey lines represent the chance level, corresponding to the probability of a random action among the 8 possible options.

Our results indicate that within each trajectory segment, the Goal-Directed model corresponding to the correct goal (e.g., Goal 1 in segment 1) exhibits the highest likelihood, with the second-based model being most of the times the Visibility-based model. Conversely, the model representing the previously achieved goal consistently shows the lowest likelihood, suggesting participants are least likely to revert to the goal they have just reached. To determine the significance of differences in likelihood values, we performed a one-way ANOVA for each condition shown in Figure 6. The model effect is significant across all conditions.

Furthermore, the pairwise comparisons (Tukey HSD) rejected the null hypothesis (*p*_*value*_ ≪ 10^−^3) in all cases except for GD3 and GD4 in Goal 2, for levels with four goals (*Mean*_*Diff*_ = 0.0004, *p*_*value*_ = 0.06), see Table S1.

Furthermore, a control analysis shows that the Goal-Directed model corresponding to the correct goal outperforms the other Goal-Directed models also during the first ten steps of each segment, permitting to rule out the possibility that its advantage is due to late phases when participants see the goal (subsection A.2).

#### 3.1.2 Participants’ Goal-Directedness is influenced by structural map characteristics

Having assessed that participants generally follow goal-directed strategies, we next asked whether this choice of strategy is fixed or modulated by map characteristics, e.g., how visible the goal is during navigation. For this, we considered whether participants’ Relative Goal-Directedness (estimated for each participant and map used the methods described in subsubsection 2.3.5) is modulated by two map characteristics:

- Goal Exposedness: This score measures the visibility of a goal from the remaining states of the map. A value of 1 indicates that the goal is fully visible from every location of the map without any obstruction, whereas a value of 0 indicates the opposite.
- Goal Online Disclosure: This metric represents the moment in the trial when the goal becomes visible to the participant. The values 0, 0.5 and 1 indicate that the goal becomes visible at the beginning of the path to the next goal, halfway during the path, or at the end of the path, respectively.

We specified two linear mixed-effects models, each with a random intercept per subject, to evaluate how Relative Goal-Directedness is associated by Goal Exposedness and Goal Online Disclosure, respectively. In both models, Relative Goal-Directedness served as the outcome variable, while Goal Exposedness and Goal Online Disclosure were entered as fixed-effect predictors in the first and second analyses, respectively. Our results reveal that Relative Goal-Directedness is positively associated with Goal Exposedness (*β* = 1.12, *t*(1.75 *×* 10^5^) = 65.20, *p <* .001), and negatively associated with Goal Online Disclosure (*β* = −.62, *t*(1.75 *×* 10^5^) = −45.48, *p <* .001)

These results imply that the relative weight of the Goal-Directed strategy increases when goals are more visible from a participant’s position and when they are revealed earlier in the trial. However, importantly, the Goal-Directed strategy has a greater weight compared to the Visibility-based strategy even in cases when goals are partially or totally obstructed (i.e., with low values of Goal Exposedness) and when they are revealed late in the trial (i.e., with high values of Goal Online Disclosure). This can be appreciated by the fact that most data points in Figure 7 lie above the dotted line, indicating a positive Relative Goal-Directedness. These results permit ruling out the possibility that participants only follow goal-directed strategies when they observe the goals.

**Figure 7:**
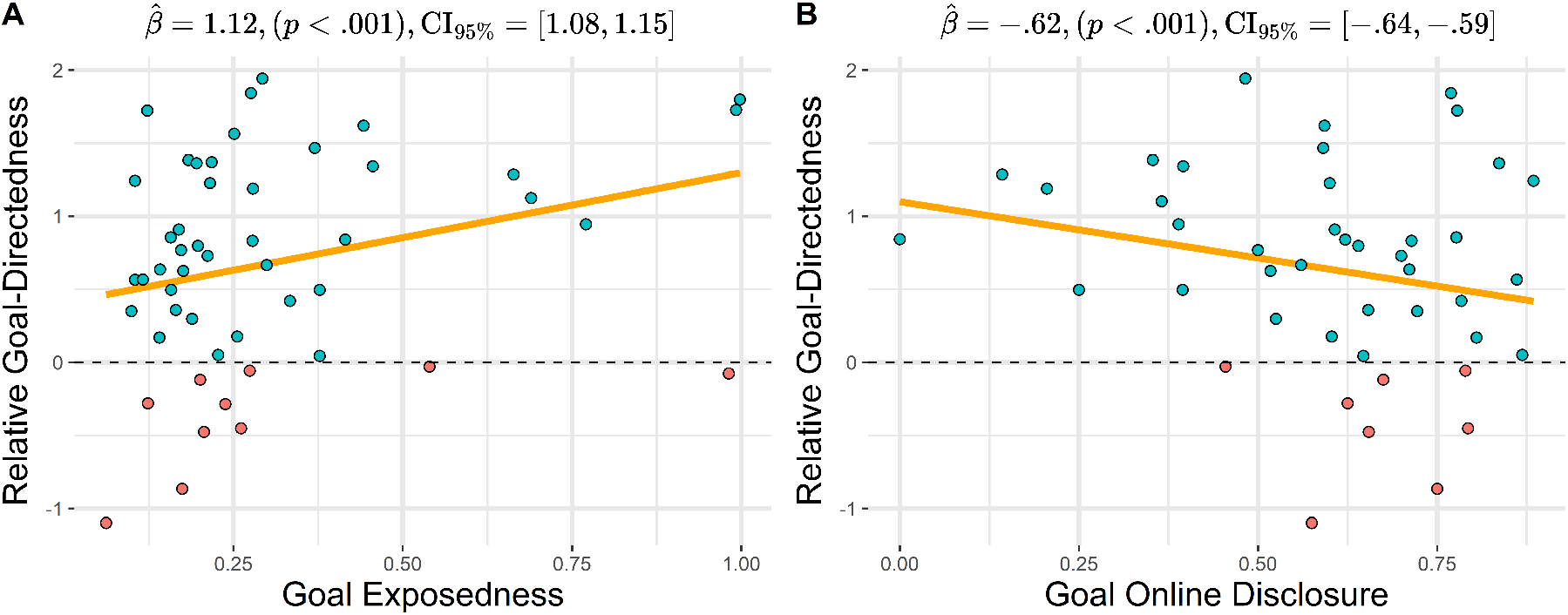
Influence of Goal Exposedness and Goal Online Disclosure on Relative Goal-Directedness. A) Scatter plot illustrating the relationship between Relative Goal-Directedness and Goal Exposedness. Each dot represents the average Relative Goal-Directedness conditioned on every Goal Exposedness score computed for every goal across all maps. B) Scatter plot depicting the relationship between Relative Goal-Directedness and Goal Online Disclosure. Each dot represents the average Relative Goal-Directedness conditioned on every Goal Online Disclosure score computed for every goal across all maps. In both figures, red and blue dots highlight values of Relative Goal-Directedness below and above the zero value, respectively.

### 3.2 Participants show signatures of sequential planning both when they memorize the map and during navigation

Having assessed that participants show goal-directed navigation, we next asked whether we could identify signatures of sequential planning, both when participants memorize the problem maps and during their subsequent navigation. We did this in two ways. First, we investigated whether participants’ view time during map memorization reflects the complexity of the problem to be solved, over and above perceptual characteristics of the map, such as its size. Second, we investigated whether during navigation we could identify signatures of sequential memory effects. We describe these two analyses in order.

#### 3.2.1 Participants’ view time reflects problem difficulty during navigation, above and beyond map size

We reasoned that if participants take into account the difficulty of solving the levels during the memorization phase, their view time (i.e., the time they spend looking at the map during the memorization phase) should be correlated with level navigation time – reflecting a greater (smaller) cognitive effort required to memorize a suitable navigation plan for difficult (easy) levels (Lancia et al., 2023; Todorov, 2009; Rubin et al., 2012b; Zenon et al., 2019). Also, if the difficulty of a map is not fully explained by trivial properties of the map, this correlation should be greater than the one between view time and simple perceptual characteristics, such as the area of the map that can be navigated.

Figure 8A plots the relations between map view time (when participants are asked to memorize the map, before navigation) and the navigation time required to solve each level, averaged over all participants (see also subsection A.4 for additional analyses). The levels are labeled with the same numbers indicated in Figure 2, for ease of comparison. The results show a significant positive correlation between the two variables (Pearson’s *r* = 0.861), indicating that levels requiring more time to view the map also tend to have longer navigation times. The correlation between map view time and the area of movable space for each map is much weaker (Pearson’s *r* = 0.358), indicating that map view time is better correlated with the difficulty to solve the problem than with simple visual properties of the map such as its size. Relatedly, follow-up analyses indicate that map view time also correlates with two other indexes of level difficulty: map tortuosity and map exposedness (subsection A.5). Taken together, these results suggests that participants take into account the (expected) difficulty of the level during the memorization phase and spend more (or less) time forming a navigation plan (as reflected by view time) for the difficult (easy) levels (i.e., as indexed by average navigation time).

**Figure 8:**
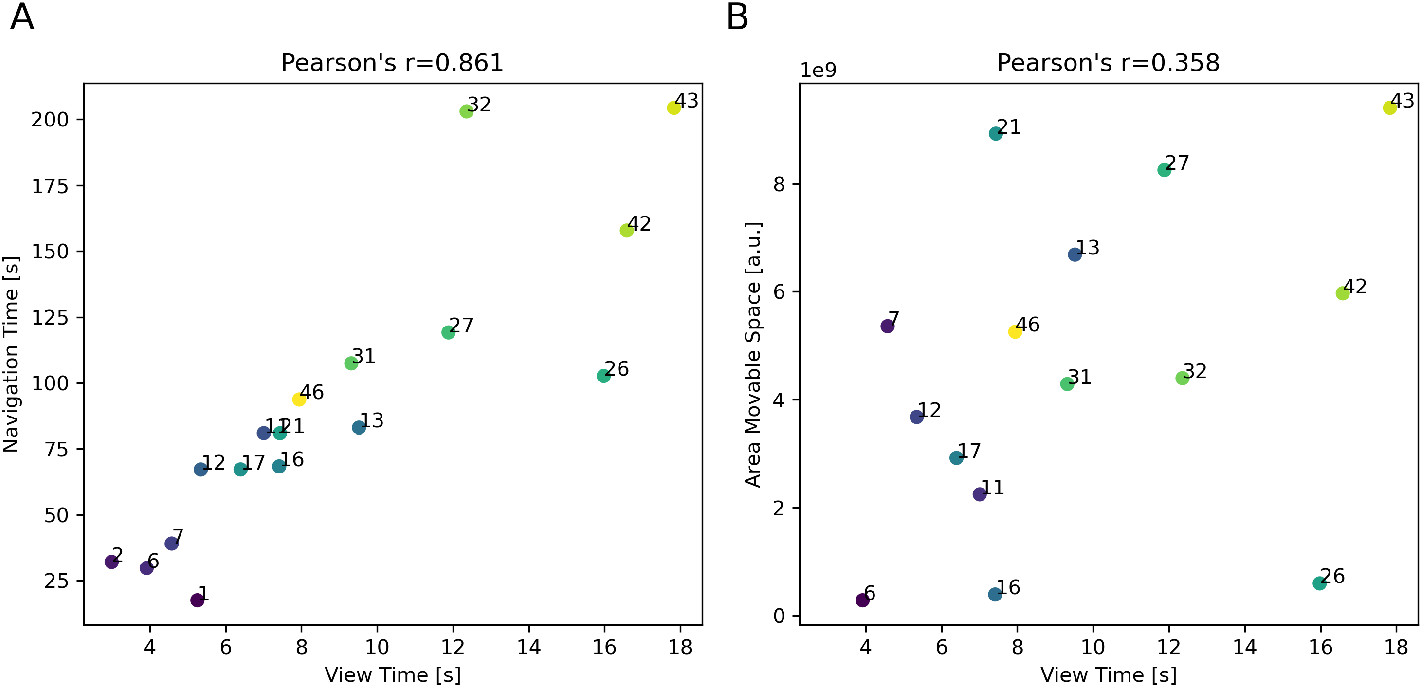
View time of SHQ levels shows high correlation with navigation time and limited correlation with map size. Scatter plots illustrating the relationship between average map view time and average map navigation time (A) or map size (B) for all the SQL levels considered in this study, with each point representing a distinct level. Data are averaged over all the participants. The levels are labeled numerically, using the same numbers as in Figure 2 for ease of comparison. The Pearson correlation coefficient is much greater for map navigation time (*r* = 0.861) compared to map size (*r* = 0.358) suggesting that map view time correlates much more with the time required to navigate the map compared to mere map size. The colors of the points represent different levels, with lighter colors indicating higher values for both metrics.

#### 3.2.2 Participants show signatures of sequential memory during navigation

Memory holds a central role in navigation, and it is known to affect the way people navigate in virtual environments (Tuena et al., 2020). The role of memory is especially important in SHQ, which requires remembering a sequence of goal locations. We reasoned that if participants form sequential navigation plans, they should show a *primacy* effect, reflecting a loss of capacity to remember goals that come later in the sequence (Ebbinghaus, 2013; Steiner and Rain, 1989; Crano, 1977; Laczó et al., 2021; Hilton et al., 2021, 2023). In this case, they should show a linearly worse performance when reaching subsequent goals. However, this effect might be mitigated by the fact that during navigation participants get exposed to the goals that come later in the sequence and could remember their position.

##### Primacy effect in navigation time

We first asked whether participants’ navigation time across goals showed a primacy effect (i.e., shorter navigation time to reach the first goal). To address this question, we analyzed the time *t* it took for participants to reach each goal from the previous (or from the start in case of Goal 1) separately for maps with 3 and 4 goals. We then compared it to the least time *t*^∗^ it would take to reach the goal by following the shortest path. From these data, we calculated the normalized time *t/t*^∗^, which is a measure of how much longer it takes for participants to reach a goal compared to the shortest path. We found that in levels with both 3 goals (Figure 9A) and 4 goals (Figure 9B), the median navigation time is smaller for Goal 1 compared to Goals 2 and 3, reflecting a primacy effect – or a better memory for the first goal in the sequence. Furthermore, in levels with 4 goals, the median navigation time is smaller for Goal 4 than for the other goals, possibly reflecting the fact that participants got exposed to the fourth goal while navigating to the other goals.

**Figure 9:**
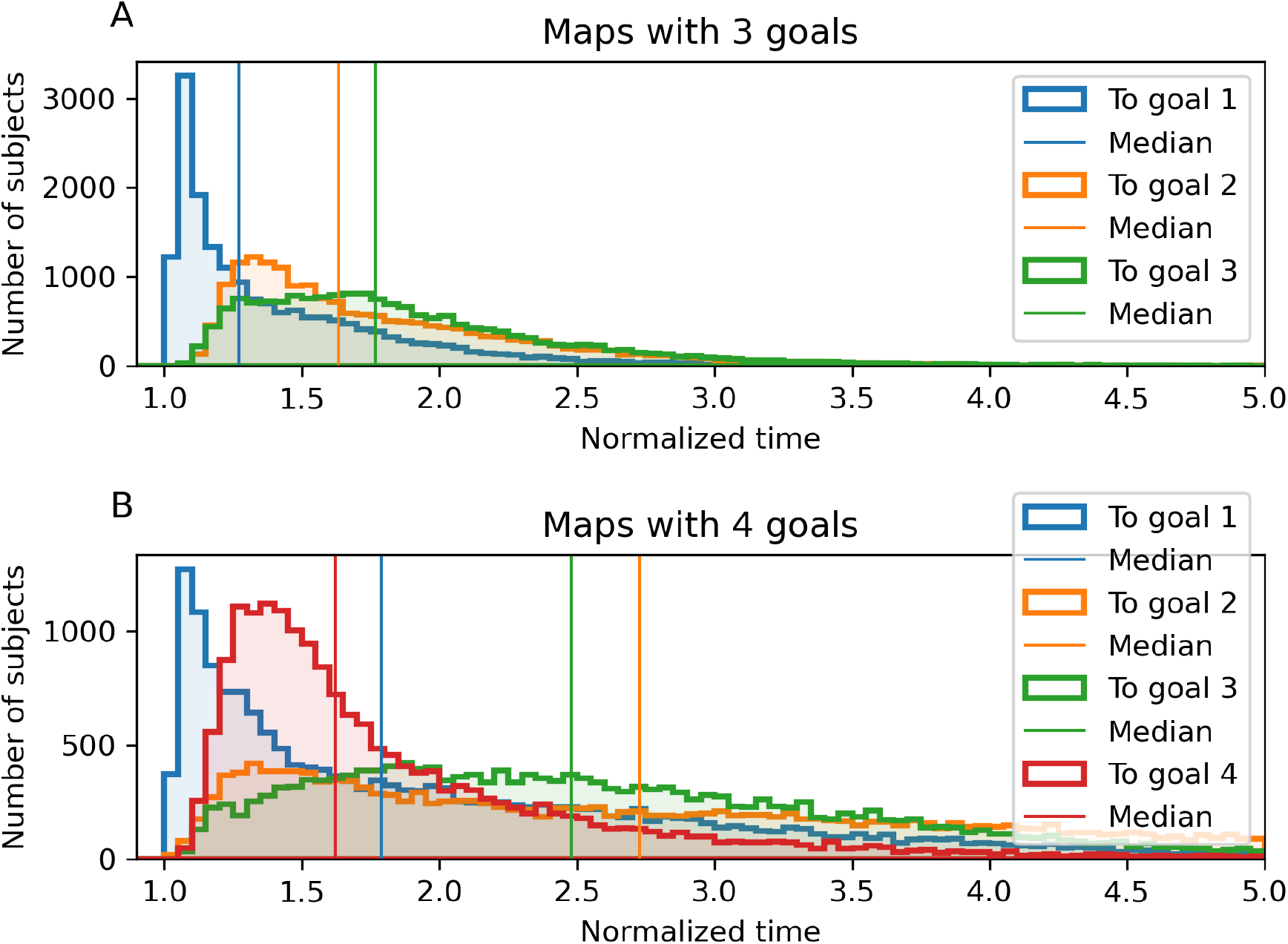
Primacy effect in navigation time. The histograms show the normalized times to arrive to Goal 1 from Start (blue), from Goal 1 to Goal 2 (orange), from Goal 2 to Goal 3 (green) and from Goal 3 to Goal 4 (red), for levels with 3 (A) and 4 (B) goals. Vertical bars show the medians of the histograms.

##### Primacy effect in the model-based analysis

Mirroring observations about navigation time, a primacy effect is also apparent when considering the averaged likelihoods of four distinct models across participants, levels, and trajectory segments (Figure 6; see also Figure S1 for another illustration of the same data). Specifically, a decreasing trend in the likelihood of Goal-Directed models is noted as goals become sequentially distant from the start, suggesting a primacy effect. Furthermore, in levels incorporating 4 goals, the likelihood of reaching Goal 4 surpasses that for Goal 3, suggesting the possibility that as participants get exposed to the fourth goal during navigation and remember its position.

### 3.3 Navigational performance decreases with age as an effect of the decline of both motor skill and goal-directedness

Age has been shown to affect navigation in real (Ham and Claessen, 2020; Fricke et al., 2022; Colombo et al., 2017; Lester et al., 2017; Zhang et al., 2021) and virtual environments (West et al., 2023), in particular in the context of spatial memory and wayfinding. Here, we aim to assess whether an age-dependent decrease of performance is observed in SHQ. Furthermore, and most crucially, we aim to assess whether the age-dependent decrease of performance is due to a decline of motor skill (e.g., the ability to control the boat dexterously during the game), of cognitive strategy (e.g., remembering the correct goal and its location, prioritizing the next goal versus map affordances) or a combination of both factors. Distinguishing motor-related and cognitive determinants of age-dependent decrease of performance is particularly challenging and for this, we adopt a combined strategy that considers both behavioral measures (navigation time, speed and angular velocity) and our computational models providing a measure of goal-directedness.

#### 3.3.1 Effects of age on navigation time, speed, angular velocity, and relative goal-directedness

We found that navigation time, averaged across all levels, significantly increases with age 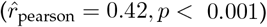 (Figure 10). Since navigation time is directly related to the number of “stars” earned during the game, this result implies an age-dependent decrease in performance. Furthermore, we found that participants’ average speed decreases with age 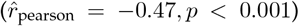, while average angular velocity, defined as the absolute value of the change in direction between consecutive steps increases with age 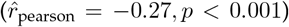. These results indicate an age-dependent decline in motor skills, suggesting that this decline is at least one of the factors contributing to the age-related increase in navigation time.

**Figure 10:**
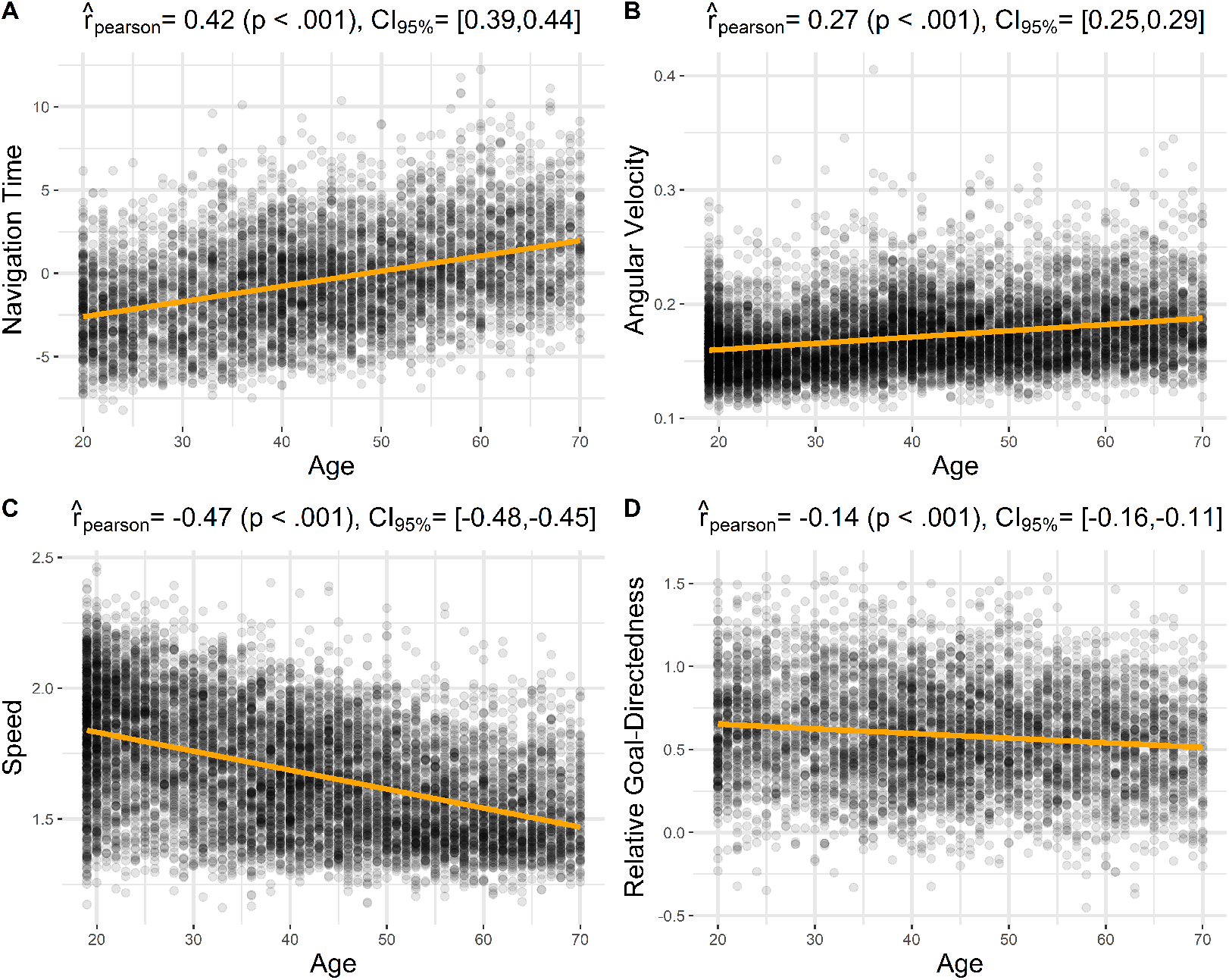
Effects of age on navigational performance. Scatter plots of (A) average navigation time of players vs. age, (B) angular velocity of players vs. age, (C) average speed of players vs. age, (D) relative goal-directedness vs. age, averaged across all levels. See the main text for explanation.

Additionally, we found that relative goal-directedness, averaged across all levels, significantly decreases with age 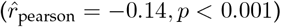. A follow-up analysis shows that this age-dependent decline in goal-directedness is also evident when considering how Goal Exposedness and Goal Online Disclosure influence Relative Goal-Directedness (subsection A.6). The fact that not only the variables indexing motor skill (angular velocity and average speed), but also our model-based measure of goal-directedness, change with age raises the question of whether motor-related and cognitive factors of performance are separable—an issue we address next.

#### 3.3.2 Separability of motor and cognitive components in navigation performance

To assess the independent contributions of motor and cognitive components to navigation performance, we aimed to demonstrate their separability in predicting participants’ behavior. For this, we used angular velocity as a proxy for motor performance and relative goal-directedness as a proxy for cognitive ability. Navigation time was used as the outcome variable, serving as a proxy for navigation performance.

We first assessed the extent to which these variables are correlated with one another. The results of the correlation analyses are illustrated in Figure 11, where each dot represents an individual participant. To better relate this analysis to age-dependent effects, we grouped participants into three distinct age cohorts, each color-coded differently: 19–35 (blue), 36–55 (orange), and 56–70 (green) years. Our analyses show that across participants, angular velocity (as a proxy for motor inefficiency) and relative goal-directedness (as a proxy for cognitive ability) relate to navigation time in opposite ways: greater angular velocity is associated with longer navigation times (Figure 11A), while higher relative goal-directedness is linked to shorter navigation times (Figure 11B). Furthermore, angular velocity and relative goal-directedness are themselves correlated (Figure 11C). These results hold across all three age groups (see Figure 11 for detailed statistical analyses).

**Figure 11:**
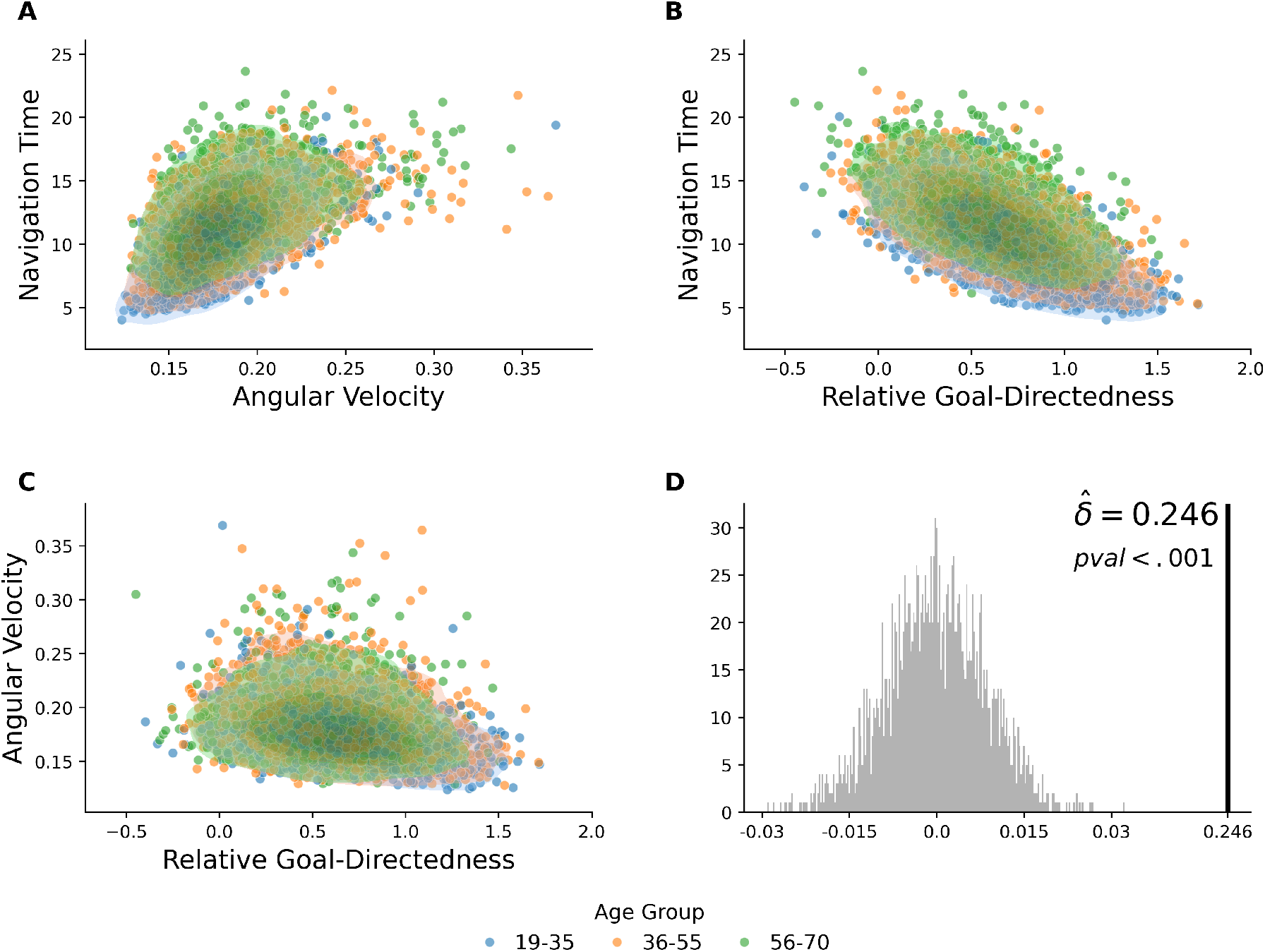
Cognitive and motor contributions to navigation performance. The three scatter plots in Panels A-C show the relationships between (A) Angular Velocity and Navigation Time, (B) Relative Goal-Directedness and Navigation Time, and (C) Relative Goal-Directedness and Angular Velocity. Dots represent individual participants and are color-coded by age cluster (blue for 19-35, orange for 36-55, and green for 56-70). The contour lines illustrate the density distribution of the data points for each age cluster. (A) Angular velocity and navigation time exhibit a positive association: 19–35 years, *ρ* = 0.617 [0.587, 0.645]; 36–55 years, *ρ* = 0.511 [0.483, 0.538]; 56–70 years, *ρ* = 0.460 [0.417, 0.500]. (B) Relative goal-directedness and navigation time display a strong negative link: 19–35 years, *ρ* = −0.680 [−0.704, −0.654]; 36–55 years, *ρ* = −0.624 [−0.656, −0.588]; 56–70 years, *ρ* = −0.545 [−0.581, −0.507]. (C) Relative goal-directedness and angular velocity show a modest inverse relationship: 19–35 years, *ρ* = −0.358 [−0.398, −0.317]; 36–55 years, *ρ* = −0.250 [−0.285, −0.214]; 56–70 years, *ρ* = −0.132 [−0.183, −0.080]. Finally, panel (D) presents the results of a non-parametric test comparing a joint model (including both cognitive and motor measures) against models using one variable alone. The histogram depicts the bootstrap null distribution.

Next, we assessed whether—despite their correlation—angular velocity and goal-directedness make independent contributions to predicting navigation time. To this end, we aggregated participants’ data by averaging angular velocity, goal-directedness, and navigation time across all goals within each map, and then across maps. We used a non-parametric bootstrap test to evaluate model improvement when both predictors are included. Specifically, we fit two models: a single-variable linear model using angular velocity to predict navigation time, and a joint model including both angular velocity and relative goal-directedness as predictors. We then computed the *R*^2^ for each model. The difference between the two, denoted as 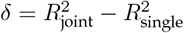, quantifies the gain in explanatory power from including the cognitive variable.

To assess the significance of *δ*, we constructed a null distribution via bootstrapping. We resampled the three variables (angular velocity, goal-directedness, navigation time) with replacement, breaking any true dependency structure, and computed the test statistic for each resample. The null distribution was centered around zero, reflecting the hypothesis of no added predictive value. The observed statistic 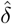 was then compared to this null distribution. As shown in Figure 11D, the observed value 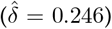 indicates a significant gain in predictive power from the joint model.

These results indicate that both motor components (angular velocity) and cognitive components (relative goal-directedness) contribute uniquely to navigation performance and do not merely reflect overlapping variance.

#### 3.3.3 Exploring the potential reasons for the decline of goal-directedness with age

Our previous analyses showed that a loss of goal-directedness contributes to the age-dependent increase of navigation time, separately from motor skill. Here, we ask whether a loss of goal-directedness might be due to the loss of capability to reach the current goal, a poor memory of the order of goals, or a shift from Goal-Directed to Visibility-Based strategies.

To address these questions, we investigated the performance of our navigational models using the same age groups (19-35, 35-55, and 55-70 years) as in the previous analyses.

In keeping with the previous analyses, for all models and trajectory segments, we observe a decreasing trend in the likelihoods of the models (across all the trajectory segments) as age increases, suggesting that older participants are less likely to follow the models’ predictions (Figure 12).

**Figure 12:**
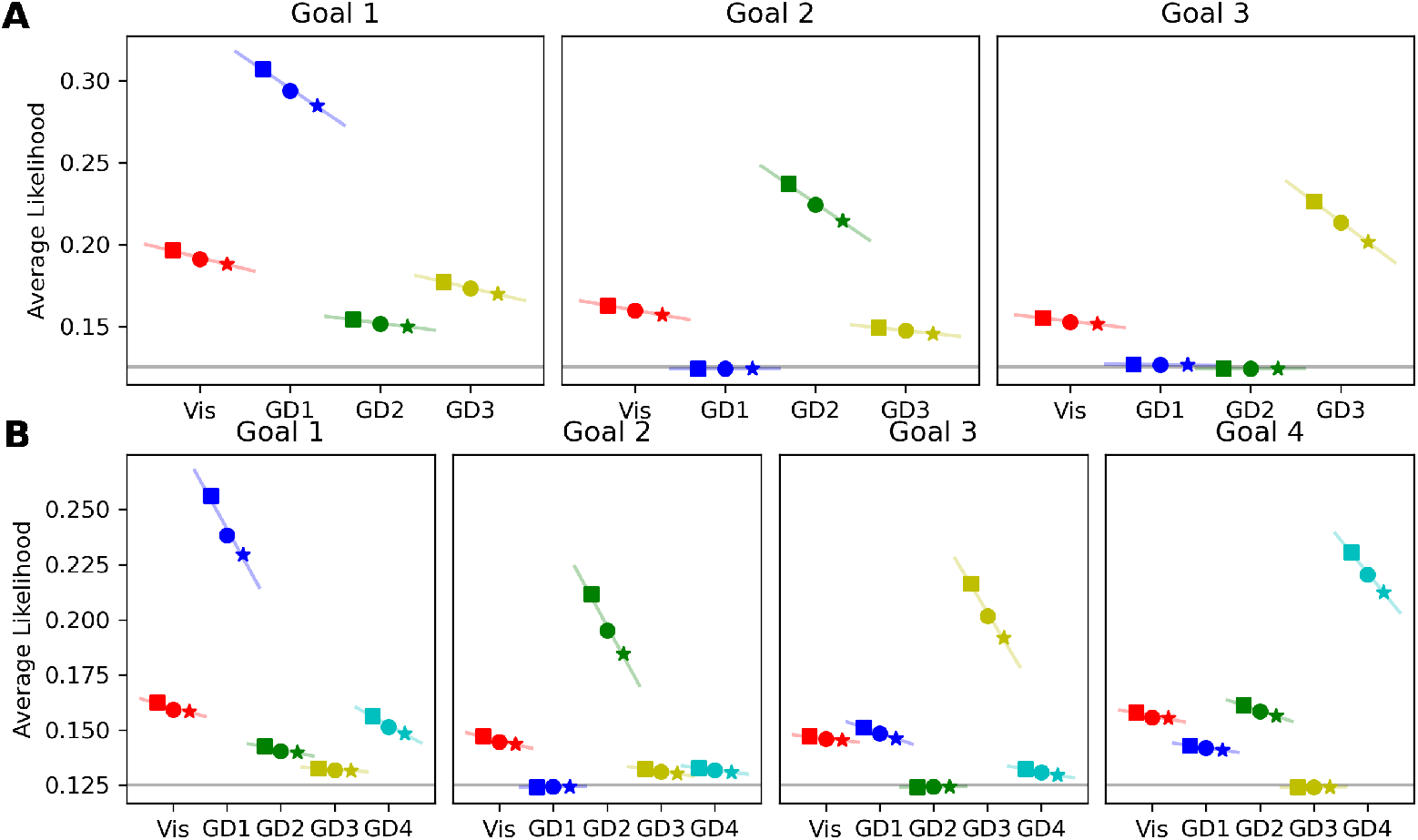
Results of the model-based analysis, illustrating the differences between age cohorts. The plots show the average likelihoods of the models as in Figure 6, but grouped by age: square markers for participants aged 19-35, circle markers for participants aged 36-55, and star markers for participants aged 56-70. The likelihoods are calculated in the same manner as previously described, with trajectory segments defined by goal reaching within each level. The models remain color-coded for ease of reference: Visibility (red), Goal-Directed model to Goal 1 (blue), Goal 2 (green), Goal 3 (yellow), and Goal 4 (azure). See Table 1 for each linear regression slope.

However, the likelihood of the “correct” goal-directed models in each trajectory segment (e.g., the Goal-Directed model to Goal 1, during the first segment) is the one that decreases the most, as indicated by the greater values of the linear regression slopes (Table 1). This result further supports the idea that the age-related decline of goal-directedness is not simply due to poorer motor skill; in such case, we should have observed a homogeneous decline of the likelihoods across all the models (as if we had added “noise” to each of them).

**Table 1:** Linear regression coefficients of slopes of Figure 12. The analysis considers all the models (Vis.: Visibility-based model; GD1-GD4: Goal-Directed models 1 to 4) and for each segment (Goal 1: from Start to Goal 1; Goal 2: from Goal 1 to Goal 2; etc.) The highest values for each segment are marked in bold.

|  |  | Vis | GD1 | GD2 | GD3 | GD4 |
| --- | --- | --- | --- | --- | --- | --- |
| Levels with 3 Goals | Goal 1 | -0.0136 | **-0.0373** | -0.0070 | -0.0127 |  |
|  | Goal 2 | -0.0096 | 0.0002 | **-0.0379** | -0.0058 |  |
|  | Goal 3 | -0.0064 | -0.0005 | 0.0002 | **-0.0413** |  |
| Levels with 4 Goals | Goal 1 | -0.0067 | **-0.0441** | -0.0046 | -0.0017 | -0.0134 |
|  | Goal 2 | -0.0058 | 0.0003 | **-0.0450** | -0.0036 | -0.0032 |
|  | Goal 3 | -0.0028 | -0.0083 | 0.0002 | **-0.0409** | -0.0044 |
|  | Goal 4 | -0.0045 | -0.0035 | -0.0081 | 0.0002 | **-0.0299** |

Furthermore, and interestingly, the decrease of the likelihood of the Goal-Directed models with age is not accompanied by an increase of the likelihood of the Visibility-based model, as one might have expected if older participants shifter from goal-directed to visibility-based strategies. Finally, a control analysis shows that a different type of cognitive error – namely, forgetting the correct order of goals – cannot explain our data (Section A.3).

Taken together, these results show that neither a decrease of motor skill, nor the passage to a (plausibly simpler) visibility-based strategy or a poor memory for the order of goals explain the age-related decline of our model-based measures of goal-directedness. While these results are not conclusive, they suggest that elder participants might lose part of their ability to reach the current goal, requiring locating themselves and/or the goal in the map, above and beyond the ability of skillful navigation.

## 4 Discussion

Spatial navigation is a fundamental and ubiquitous skill but we still have an incomplete understanding of how people make navigation decisions in real life conditions. In this study, we aimed to investigate the strategies that people adopt to solve navigation problems, using a computational approach and the rich navigational setup provided by the large scale Sea Hero Quest (SHQ) game (Spiers et al., 2023; Coutrot et al., 2019, 2018a; West et al., 2023).

Through a combination of behavioral and model-based analyses, we report three main findings. First, participants generally follow goal-directed navigation, as evidenced by the higher likelihood of the Goal-Directed models in predicting participants’ actions compared to the Visibility-based model. This occurs also in nontrivial cases such as when the goal is not visible. This finding is consistent with the idea that individuals often navigate by identifying and following the most efficient path to a predetermined goal. However, when deviating from goal-directed strategies, participants tend to follow the Visibility-based strategy rather than pursuing one of the incorrect goals. This can be appreciated by considering that the likelihood of the Visibility-based model is generally greater than the likelihoods of the incorrect Goal-Directed models (Figure 12).

Second, participants show key signatures of sequential planning. During the memorization phase, they spend more time viewing the most difficult problems (those requiring longer navigation time), possibly reflecting a greater cognitive effort to encode a more complex plan, over and above the effect of mere map size. Furthermore, during navigation, participants show a better memory for goals that come earlier, aligning well with *primacy* effects in memory for sequences (Ebbinghaus, 2013; Steiner and Rain, 1989; Crano, 1977) including studies during virtual navigation (Laczó et al., 2021). In problems with four goals, they took less time to reach the final goal, plausibly reflecting the fact that they could see such goal while navigating towards the other ones. In turn, this suggests that in embodied settings like SHQ, planning does not only occur before navigation but also during it.

Third, and most importantly, there is a significant age-dependent decrease of navigation performance, as indexed by navigation time to complete the problems. We used combined behavioral and computational measures to distinguish motor- and cognitive-related determinants of performance decrease, which are typically difficult to disambiguate. Our analyses indicate that age-dependent decrease of navigation performance could be jointly explained by a decline of both motor skill (as indexed by increased angular velocity and decreased speed of the boat) and cognitive strategy (as indexed by a decrease of our model-based indexes of goal-directed control: goal-directedness and relative goal-directedness). Our analyses reveal that, although both motor and cognitive decline are correlated with age-related decreases in navigational performance, they contribute independently to it. Unexpectedly, when deviating from goal-directed strategies, elder participants do not simply shift to visibility-based strategies, as it might be naively assumed when considering that such strategies are cognitively less demanding. Furthermore, the behavior of elder participants cannot be accounted for by considering an incorrect memory for the sequence of goals. Our analyses are therefore not conclusive about the possible sources of the age-dependent decline of goal-directedness and cognitive strategy. Such decline could encompass impaired self-localization and navigation, memory for the correct goal location, planning failures and other factors that render participant’s behavior different from the shortest-path solutions implicit in our Goal-Directed models, or other factors. The contributions of these or other factors to the cognitive decline that we observe remain to be investigated in future studies. Furthermore, future studies might assess whether the decline of cognitive strategy and spatial memory could be associated with neurodegenerative diseases, including Alzheimer’s disease.

These results are consistent with previous findings in the literature, and provide a more fine-grained understanding of the mechanisms behind the decline of spatial navigation with age. The landscape of research into decision-making and navigation strategies has been significantly enriched by the introduction of various game-based methodologies (see for example (Rafferty et al., 2012; Allen et al., 2023; Brändle et al., 2023)). Among these, SHQ ((Spiers et al., 2023)) stands out due to its large-scale global reach, offering unprecedented opportunities to study navigational decision-making across different ages, countries, and demographic groups. Data from SHQ has highlighted significant navigational ability differences across age groups, with older participants performing worse than younger ones (Spiers et al., 2023), along with variations by gender, education level, and living environment (Coutrot et al., 2018b). These findings are meaningful outside the virtual realm, correlating with real-world navigation performance (Coutrot et al., 2018b). SHQ data also suggested the game’s potential to identify genetic markers for Alzheimer’s disease (Lim et al., 2023; Coughlan et al., 2019) and demonstrated that a player’s handedness does not affect their navigation skills (Fernandez-Velasco et al., 2023).

Our investigation into SHQ not only extends the existing body of research but also introduces computational models that incorporate diverse navigation strategies, such as goal-directed navigation versus visibility-based decision-making. These models provide a sophisticated framework for analyzing the ways in which participants solve SHQ problems, allowing us to explore the potential sources of performance decline with age and (potentially) individual differences. Furthermore, the computational models developed for this study provide time-varying signals (e.g., the likelihoods of the different models in Figure 4) that could be potentially useful to analyse time series of neural data, as in electroencephalography (EEG). The methods developed in this study have therefore important implications for the study of human spatial navigation and its decline with age. By providing a more fine-grained understanding of the mechanisms behind the decline of spatial navigation with age, our work contributes to the development of more effective interventions for older adults, such as the design of training programs that target specific cognitive or motor control processes. Furthermore, our methods are general and can be applied to other video games and virtual environments, providing a new way to study human spatial navigation and its decline with age, moving towards the computational phenotyping of subjects (i.e., the description of a subjects in terms of a specific parametrization of a model). Further exploration of the mechanism related to performance decline could potentially lead to personalized interventions or training programs tailored to specific navigational strengths and weaknesses.

A potential limitation of this study is that the quantification of motor versus cognitive skills could be more nuanced than that we considered. From this perspective, certain aspects of navigation speed may reflect cognitive processing demands (e.g., managing uncertainty), rather than being solely attributed to motor control. For instance, older participants might slow down to better perceive and recognize useful map locations—although this would not easily account for their increased angular velocity. Conversely, excessive path length might be due in part to poor motor control. Future studies may consider using additional metrics to better disentangle motor versus cognitive skills. Another limitation of the study is that it does not allow full control of exposure to mobile phone games in different age groups. Furthermore, future studies might consider expanding the repertoire of computational models used in this research domain. In the present study, following a well-established literature on decision-making and navigation, we adopt the framework of Markov decision processes (Puterman, 1994; Bellman, 1958). This framework makes the simplifying assumption that the current state contains all the information the agent needs to make decisions. Moreover, we do not model cognitive map construction, nor does our model include any learning components. Alternative approaches—such as models based on reinforcement learning, probabilistic inference, or deep learning—could offer complementary perspectives on navigation strategy selection and adaptation. These models might better capture complex aspects of human navigation, such as learning from past experiences or adapting to dynamically changing environments. This is especially relevant to the formation of cognitive maps, which in our models are assumed to be known and memorized. In reality, however, such maps may be noisily encoded and dynamically updated over time. Future work could investigate how SHQ participants build cognitive maps during the initial observation phase and revise them throughout subsequent navigation. By the same token, future studies could model a broader range of possible navigation strategies (Parra-Barrero et al., 2023), beyond the goal-directed and visibility-based strategies investigated here. Comparing the predictions of these alternative models with those derived from our current approach could help clarify the relative contributions of different cognitive processes to navigational success and support the development of more comprehensive models of human spatial navigation. Finally, future research could move beyond virtual environments to investigate the extent to which the findings of this study possess ecological validity and generalize to real-world navigation tasks. This includes exploring navigation in diverse types of environments—such as urban, rural, and maritime settings—and examining how individuals navigate both familiar and novel terrains (Ekstrom et al., 2018; Maselli et al., 2023).

### Data repository

Scripts for the analysis of this paper can be freely accessed at the following repository: https://github.com/gllancia/Separable-cognitive-and-motor-decline

## Acknowledgments

This research received funding from the European Union’s Horizon 2020 Framework Programme for Research and Innovation under the Specific Grant Agreements No. 952215 (TAILOR) to G.P.; the European Research Council under the Grant Agreement No. 820213 (ThinkAhead) to G.P.; the Italian National Recovery and Resilience Plan (NRRP), M4C2, funded by the European Union – NextGenerationEU (Project IR0000011, CUP B51E22000150006, “EBRAINS-Italy”; Project PE0000013, CUP B53C22003630006, “FAIR”; Project PE0000006, CUP J33C22002970002 “MNESYS”) to G.P., PRIN PNRR P20224FESY to G.P. M. I. is supported by: the European Union under the scheme HORIZON-INFRA-2021-DEV-02-01 – Preparatory phase of new ESFRI research infrastructure projects, Grant Agreement n.101079043, “SoBigData RI PPP: SoBigData RI Preparatory Phase Project”; by the project “Reconstruction, Resilience and Recovery of Socio-Economic Networks” RECON-NET EP FAIR 005 - PE0000013 “FAIR” - PNRR M4C2 Investment 1.3, financed by the European Union – NextGenerationEU; by the European Union - Horizon 2020 Program under the scheme ‘INFRAIA-01-2018-2019 - Integrating Activities for Advanced Communities’, Grant Agreement n.871042, ‘SoBigData++: European Integrated Infrastructure for Social Mining and Big Data Analytics’ (http://www.sobigdata.eu). The GEFORCE Quadro RTX6000 and Titan GPU cards used for this research were donated by the NVIDIA Corporation. The funders had no role in study design, data collection and analysis, decision to publish, or preparation of the manuscript.

## A Supplementary materials

### A.1 Comparison between Goal-Directed and Visibility-based models – additional plots

Figure S1 and Figure S2 show alternative ways to plot the comparison between Goal-Directed and Visibility-based models, when averaging across all age cohorts and when considering age cohorts separately, respectively.

**Figure S1:**
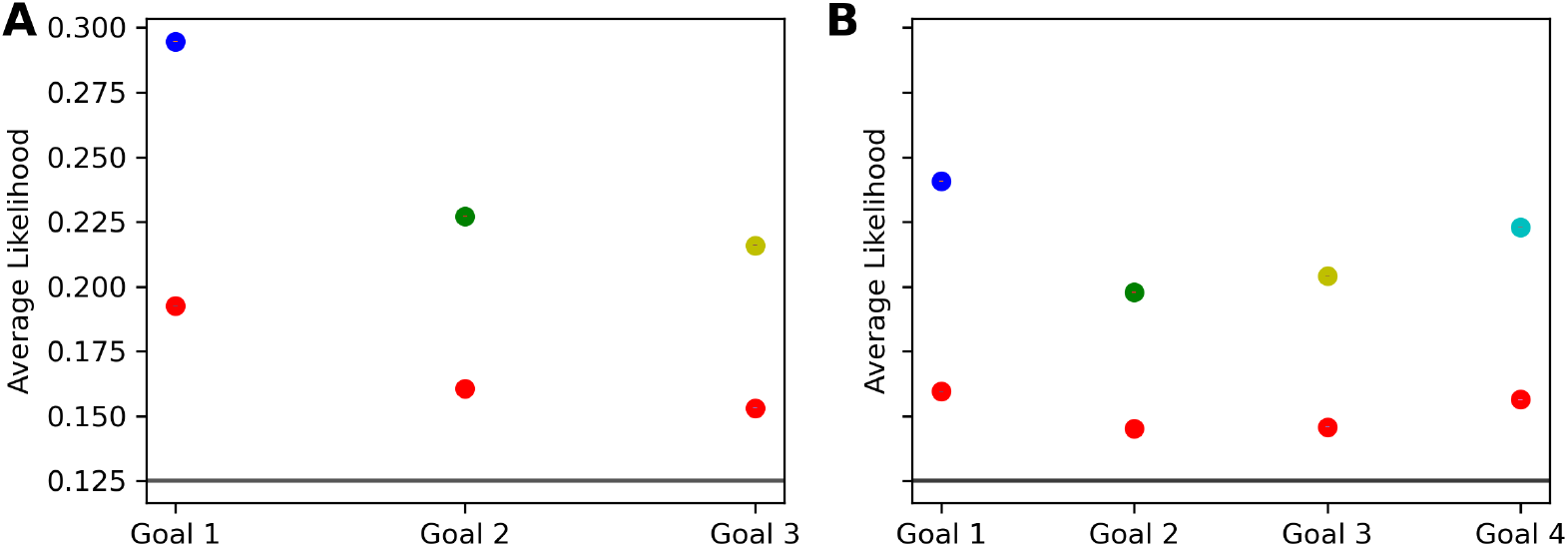
Likelihood of the relevant Goal-Directed (GD) models. Panels A and B summarize the likelihood of the relevant Goal-Directed (GD) models for levels with three and four goals, respectively, plotting only the likelihoods for the “correct” GD model for each goal (blue, green, yellow, azure) alongside the Visibility-based model (red).

**Figure S2:**
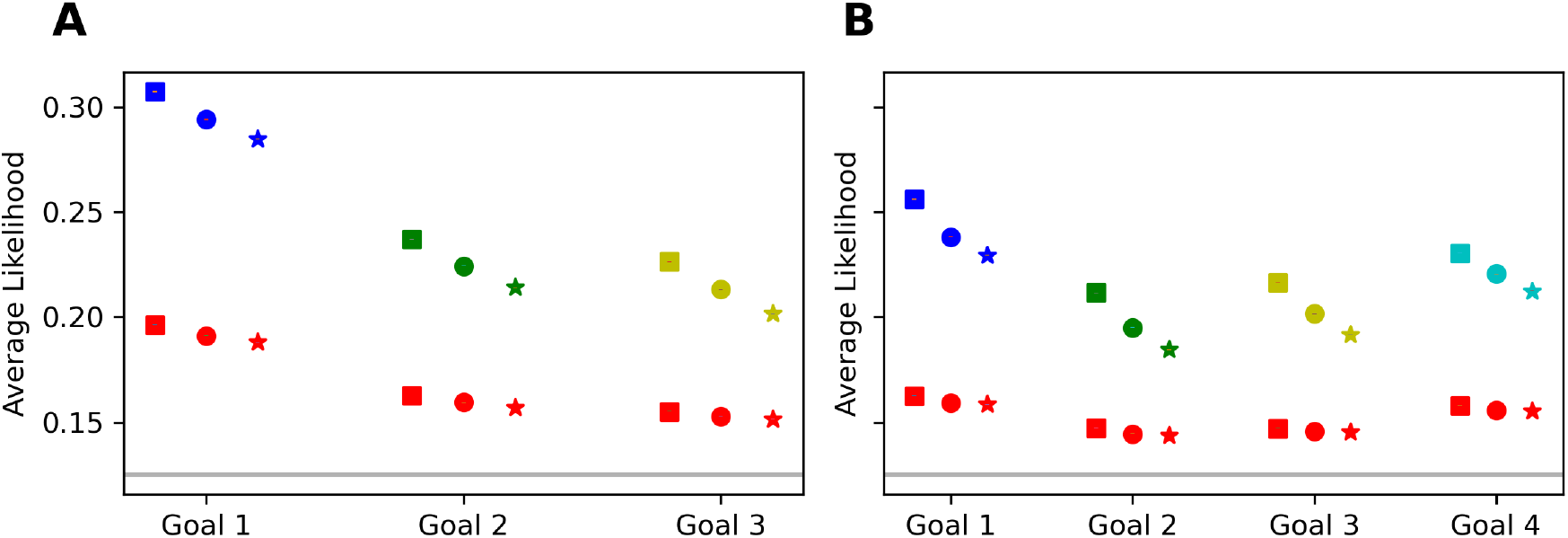
Model Performance by Goal, Level and Age Cohort. Averaged likelihoods as in Figure S1, but grouped by age: square markers for participants aged 19-35, circle markers for participants aged 35-55, and triangle markers for participants aged 55-70. The likelihoods are calculated in the same manner as previously described, with trajectory segments defined by goal reaching within each level. The models remain color-coded for ease of reference: Visibility (red), Goal-Directed model to Goal 1 (blue), Goal 2 (green), Goal 3 (yellow), and Goal 4 (azure).

### A.2 Comparison between Goal-Directed and Visibility-based models, during the first 10 navigation steps

As an additional way to compare Goal-Directed and Visibility-based models, we analysed their likelihoods in the same way as in the main text, but by only considering the initial 10 steps of participants’ trajectories across all levels. Focusing on the initial stage of the trajectory enables differentiation of the initial strategies used by participants, which are indicative of their plans.

Figure S3 shows the results of the analysis. These results are largely congruent with those shown in Figure 6) and confirm that the “correct” models (e.g., Goal-Directed model 1 for the first segment of trajectory) showing the highest likelihoods. Note that the Visibility-based model outperforms all the others in the initial trajectory segment towards Goal 1. However, this result likely depends on the mere fact that almost all starting points are from corridors or corridor-like structures, thereby enhancing the Visibility-based model’s likelihood in early participant behavior – an effect that is also apparent when inspecting our example of model-based analysis in Figure 4.

Figure S4 shows the results of the analysis, when splitting per age cohorts (as in Figure 12. Consistent with previous findings, our analysis revealed an age-related decline in model likelihoods, with “correct” models (e.g., Goal-Directed model 1 for the first segment of trajectory) showing the highest likelihoods across age groups.

**Figure S3:**
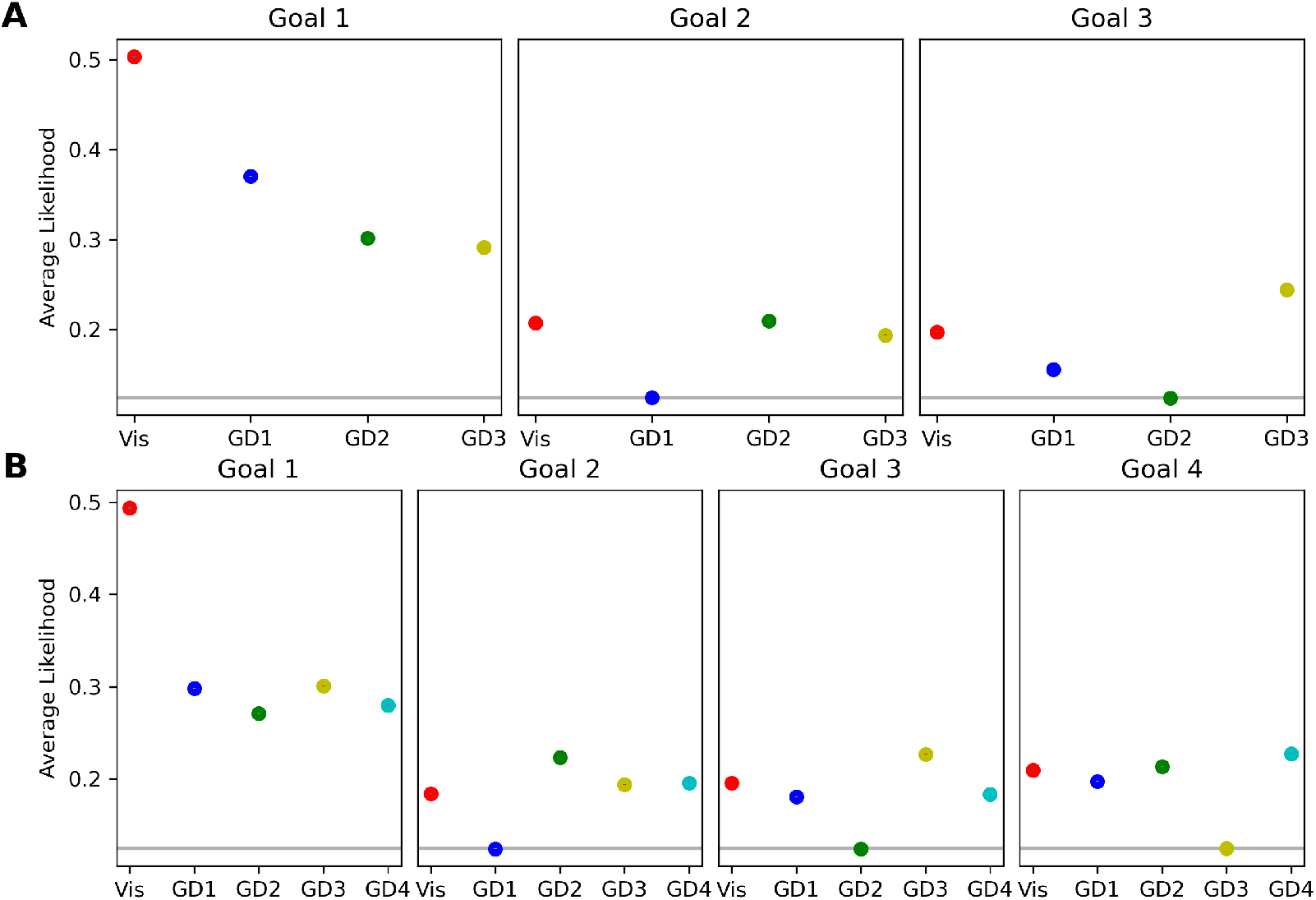
Model Performance by Goal and Level as in Figure 6, but for the first 10 steps. Averaged likelihoods of the models over participants, levels, and trajectory segments. Error bars represent standard errors, which are smaller than the depicted symbols. The models are color-coded: red for Visibility, blue for Goal-Directed to Goal 1, green for Goal 2, yellow for Goal 3, and azure for Goal 4. Panel A (top row) displays model likelihoods for levels with three goals, with subplots for each goal reflecting model performance as participants navigate towards each successive goal. Panel B (middle row) shows likelihoods for levels with four goals

**Figure S4:**
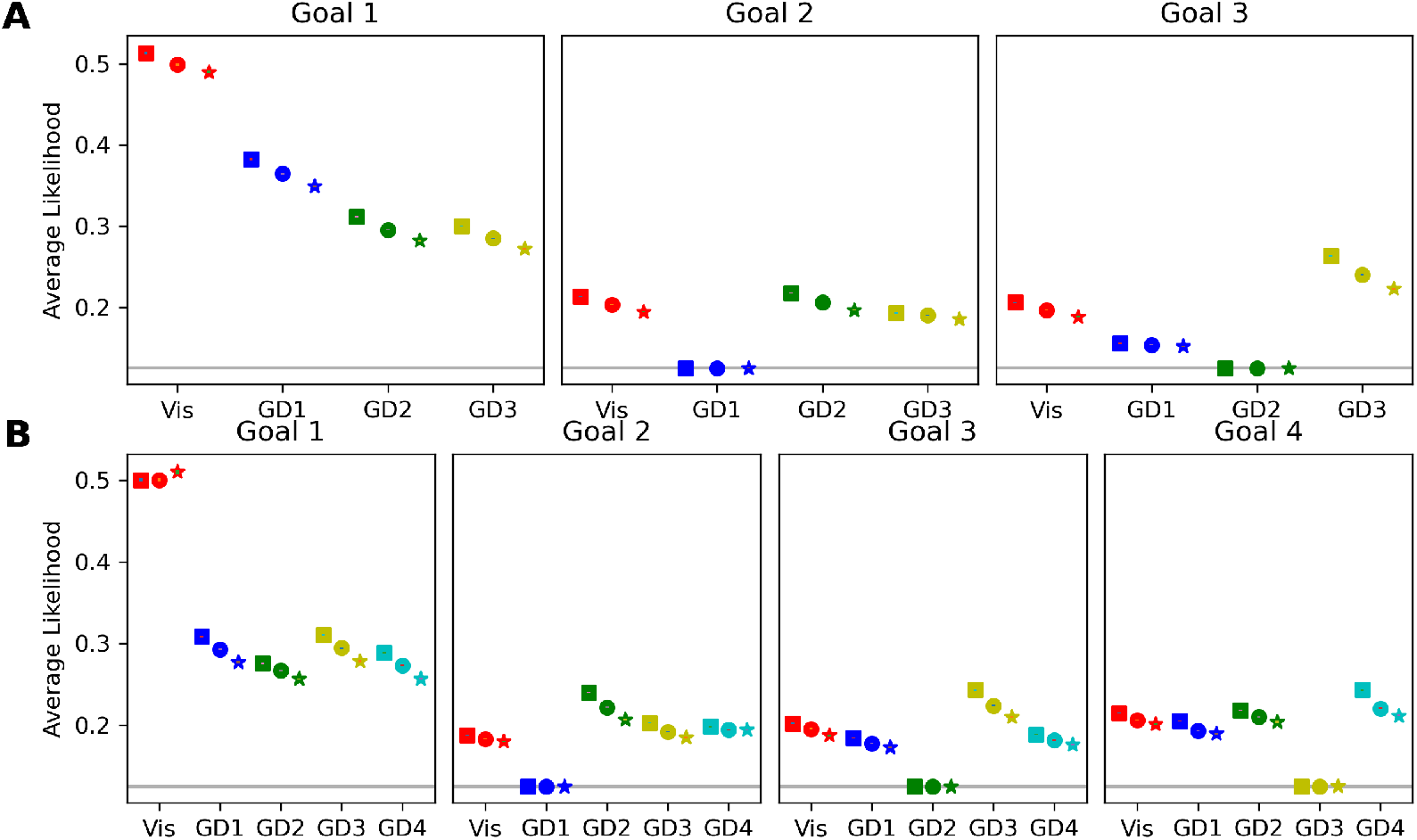
Model Performance by Goal, Level and Age Cohort as in Figure 12, but for the first 10 steps. Averaged likelihoods grouped by age: square markers for participants aged 19-35, circle markers for participants aged 35-55, and star markers for participants aged 55-70. The likelihoods are calculated in the same manner as previously described, with trajectory segments defined by goal reaching within each level.The models remain color-coded for ease of reference: Visibility (red), Goal-Directed model to Goal 1 (blue), Goal 2 (green), Goal 3 (yellow), and Goal 4 (azure

### A.3 Participants tend not to to confuse the correct order of goals during navigation

We tested the hypothesis that the decreased goal-directedness reported in elder participants could be explained by the fact that they forget the correct sequence of goals, therefore pursuing an incorrect goal.

To model this sequencing error, we introduced a novel, *Sequencing Error* model that arbitrates between all shortest path models by introducing an epsilon-greedy switching mechanism, that upon reaching any goal — correct or incorrect — selects the next one, which will be with a probability *p* the correct goal and 1 − *p* another random goal. We varied the *p* parameter to three values (*p* = 0.99, 0.8, 0.6) and analyzed the models’ likelihoods in the initial 10 steps, as we did with the participants.

The Sequencing Error model (Figure S5) showed a decrease in likelihood for the “correct” model with a decreasing *p*, while the likelihoods for other models increased. This confirms that as *p* decreases—increasing the propensity to select incorrect goals—the likelihood for models targeting the correct goal diminishes, while it increases for other models, as predicted. This effect contrasts with the experimental results (Figure 12). This suggests that the decline in performance with age cannot be explained by a lack of memory for the correct sequence of goals.

**Figure S5:**
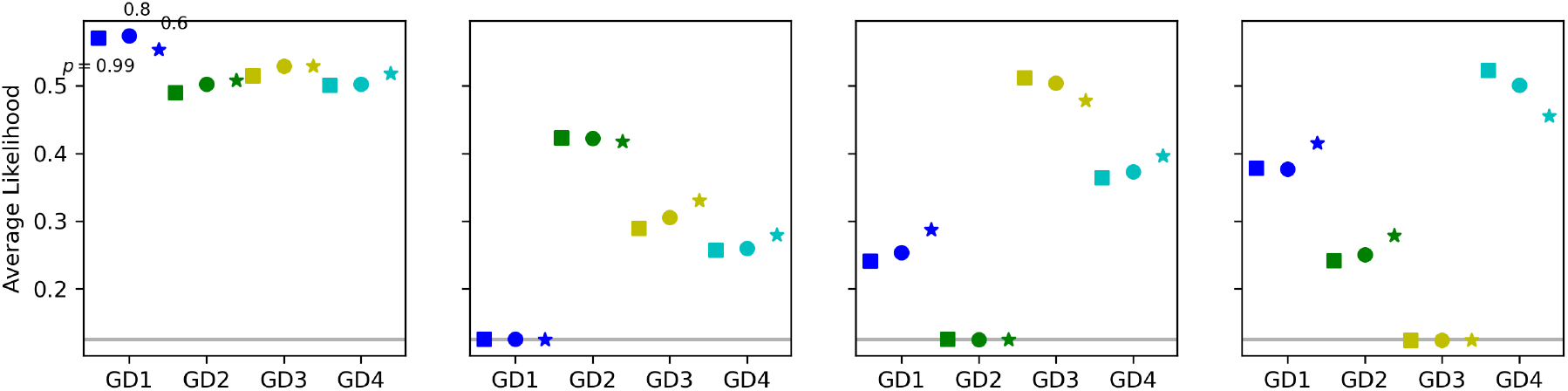
Model Performance by Goal, Level, and Age Cohort in Early Trajectory Phases with data simulated using a Sequencing Error model. The figure shows the averages of likelihoods of the Sequencing Error model’s simulated trajectories evaluated at three distinct values of *p* (*p* = {0.99, 0.8, 0.6}). For simplicity, simulation was performed for the 10 time steps of each problem, which are the most discriminative. See the main text for explanation.

### A.4 Correlation between navigation and view time within levels

We examined the relationship between navigation time and view time separately for each level, using the biweight mid-correlation: a robust estimator that recentres the data on the median, rescales it by the median absolute deviation, and progressively down-weights observations the farther they stray from the centre—eventually giving extreme points no leverage at all. Because our data exhibited occasional spikes and heteroscedastic segments, this robust treatment preserved all observations while limiting the influence of outliers; hence, it provided a principled alternative to ad-hoc outlier removal and yielded more reliable estimates of the underlying linear relationships. A correlation was deemed statistically significant when its 95% confidence interval excluded zero. (Figure S6).

Our results show that, for 10 out of 17 levels, there is a significant positive correlation between view time and navigation time, indicating that participants who spent more time viewing the level tended to navigate it more slowly. For 5 levels, we observed a significant negative correlation, suggesting that longer viewing was associated with faster navigation. Notably, 4 of these 5 levels contain 4 goals, which may indicate a more demanding planning process (the only other level with 4 goals not included in this group is level 27). Finally, for 2 levels, the correlation between view time and navigation time was not significant.

**Figure S6:**
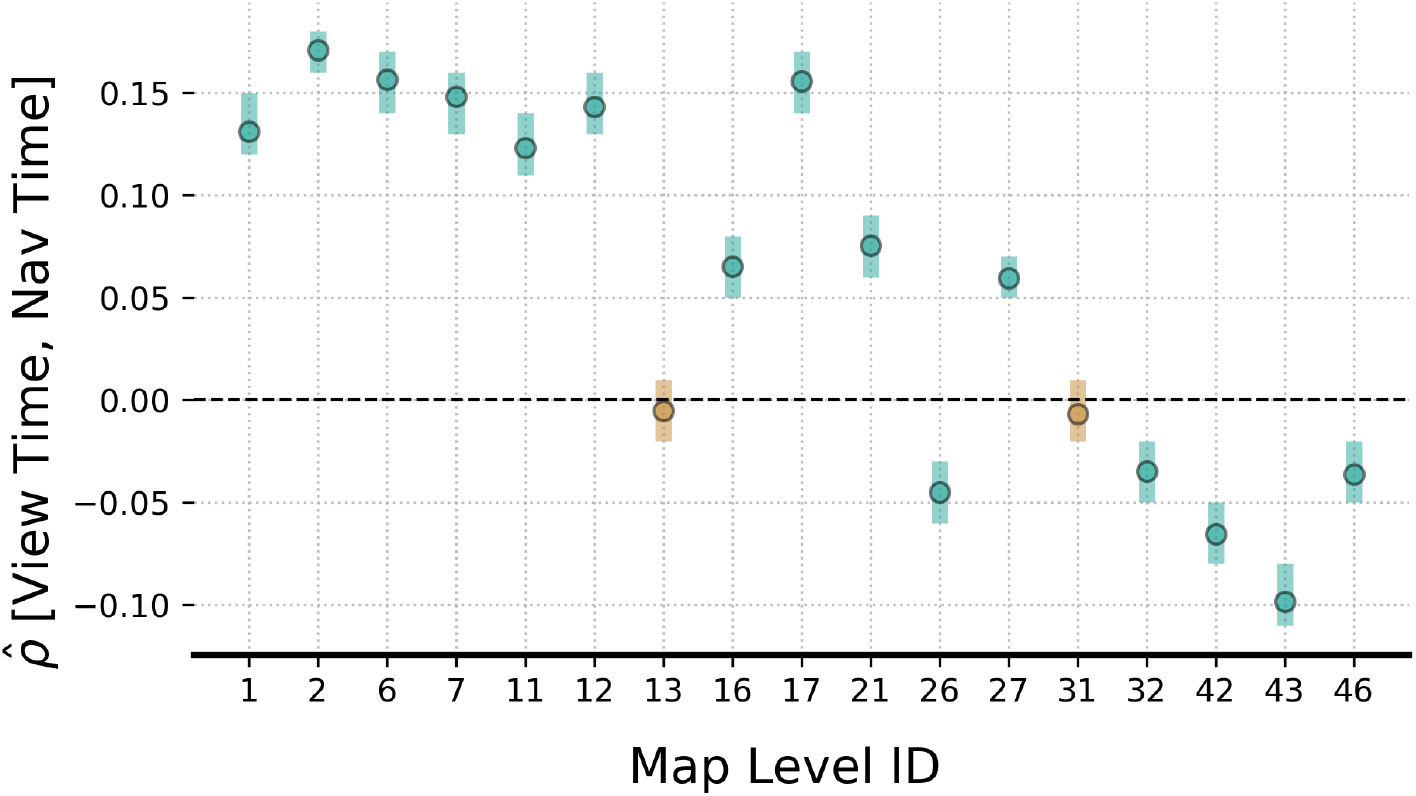
Within-map correlation analysis of map navigation time and map view time. Correlation values between navigation time and view time are depicted for every map level. Dots represent the correlation effect size with 95% confidence intervals error bars. Green and brown colors represent significant and non-significant correlation values, respectively.

### A.5 Influence of map tortuosity and map exposedness on map view time

Figure 8 shows a positive correlation between the time participants spend viewing the problem maps (possibly reflecting planning demands) and their subsequent navigation time (possibly reflecting map difficulty). We performed a follow-up analysis, to assess whether map view time correlates also with two other metrics of map difficulty:

- Map Tortuosity: This metric is computed as the average tortuosity score across all goals on the map. The tortuosity score for each goal is defined as the ratio between the actual path length from a seed (such as a starting point or a previous goal) to the goal, and the straight-line distance between them.
- Map Exposedness: This metric is computed by averaging the Goal Exposedness values across all goals on the map. Each Goal Exposedness value quantitatively measures how visible a goal is from various points on the map, with a value of 1 indicating complete visibility from everywhere on the map.

We performed two separate mixed-effect regression analyses with subjects as random effects. The map’s view time is used as the dependent variable in the two map-level analyses, and Map Tortuosity and Map Exposedness as predictors, respectively for each analysis. For all the analyses, clustered Age (19-35, 36-55, 56-70) was used as a covariate.

Results of the first analysis show a significant increase in the map’s view time as Map Tortuosity increases (*β* = 0.39, *t*(1.7 *×* 10^5^) = 90.78, *p <* 0.0001), and a significant effect of age. In particular, a contrast analysis revealed that intercepts increase for older age clusters (Figure S7, A), such that the overall map’s view times is higher for the 36-55 age cluster compared to 19-35 (*β* = 0.158, *z* = 10.97, *p <* 0.0001), and higher for 56-70 compared to 36-55 (*β* = 0.22, *z* = 14.31, *p <* 0.0001). Results of the second analysis show a significant decrease in the map’s view time as Map Exposedness increases (*β* = −1.48, *t*(1.7 *×* 10^5^) = −147.29, *p <* 0.0001), and a significant effect of age on the intercepts as well (Figure S7, B). In particular, the overall map’s view times is higher for the 36-55 age cluster compared to 19-35 (*β* = 0.16, *z* = 11.05, *p <* 0.0001), and higher for 56-70 compared to 36-55 (*β* = 0.23, *z* = 14.53, *p <* 0.0001). Altogether, these findings indicate that individuals require more time to encode maps that exhibit greater difficulty (characterized by increased tortuosity) and those that prospect more challenging navigation (indicated by lower exposedness), as well as that younger individuals are faster at encoding maps overall.

**Figure S7:**
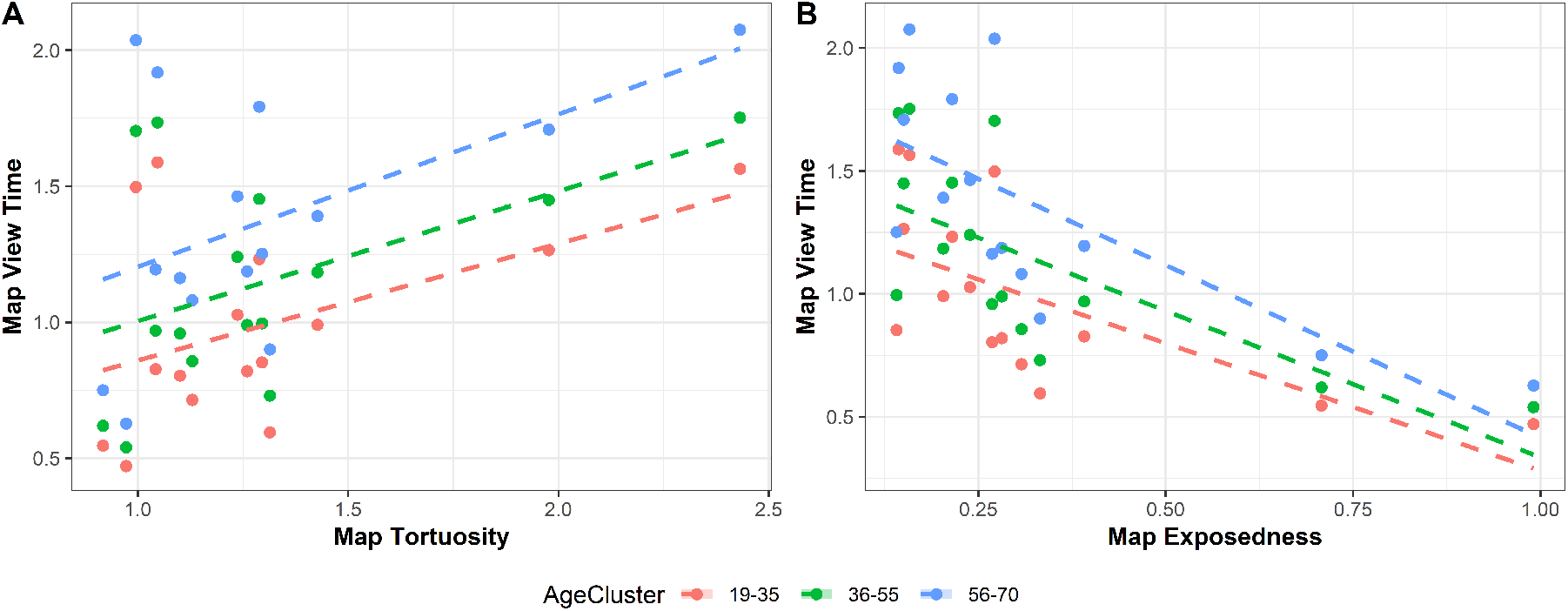
Relationship Between Map View Time and Map Characteristics Across Age Clusters. A) Scatter plot showing the relationship between map view time and Map Tortuosity (see main text). Each dot represents the value averaged across all subjects in a given level, color-coded by age cluster (red for 19-35, green for 36-55, and blue for 56-70). The dashed lines represent the linear regression fits for each age cluster, indicating an increase in map view time with increased Map Tortuosity. B) Scatter plot illustrating the relationship between map view time and Map Exposedness. Each dot represents the value averaged across all subjects in a given level, color-coded by age cluster (red for 19-35, green for 36-55, and blue for 56-70). The dashed lines show the linear regression fits for each age cluster, indicating a decrease in map view time with increased Map Exposedness.

We also tested whether there is a possible correlation between view time and goal-directedness, but the correlation does not reach significance (p *>* 0.5).

### A.6 Influence of Goal Exposedness and Goal Online Disclosure on Relative Goal-Directedness Across Age Clusters

In the main text, we found correlations between Relative Goal-Directedness and both Goal Exposedness and Goal Online Disclosure. Here, we perform a follow-up analysis to further investigate the relations between these variables and age. For this, we performed a mixed-effect regression analyses with subjects as random effects. For all the analyses, the age of participants clustered in three cohorts (19-35, 36-55, 56-70) was used as a covariate. Relative Goal-Directedness was first estimated given the model in eq. 6 for each subject and each goal, and its logit was used as the dependent variable. Goal Exposedness and Goal Online Disclosure were used separately as predictors for each analysis, respectively.

Results of the first analysis show a significant increase in Relative Goal-Directedness as Goal Exposedness increases (*β* = 1.12, *t*(1.7 *×* 10^5^) = 65.42, *p <* 0.0001) and a significant effect of age (Figure S8A) such that the overall Relative Goal-Directedness is higher for the 19-35 age cluster compared to 36-55 (*β* = 0.05, *z* = 5.85, *p <* 0.0001), which in turn shows higher values compared to 56-70 (*β* = 0.06, *z* = 6.43, *p <* 0.0001).

Results of the second analysis reveal a significant decrease in Relative Goal-Directedness as Goal Online Disclosure increases (*β* = 0.62, *t*(1.7 *×* 10^5^) = −45.61, *p <* 0.0001) and a significant effect of age on the intercept (Figure S8B). Even in this case, the overall Relative Goal-Directedness is higher for the 19-35 age cluster compared to 36-55 (*β* = 0.05, *z* = 5.55, *p <* 0.0001), which in turn shows higher values compared to 56-70 (*β* = 0.057, *z* = 5.82, *p <* 0.0001).

**Figure S8:**
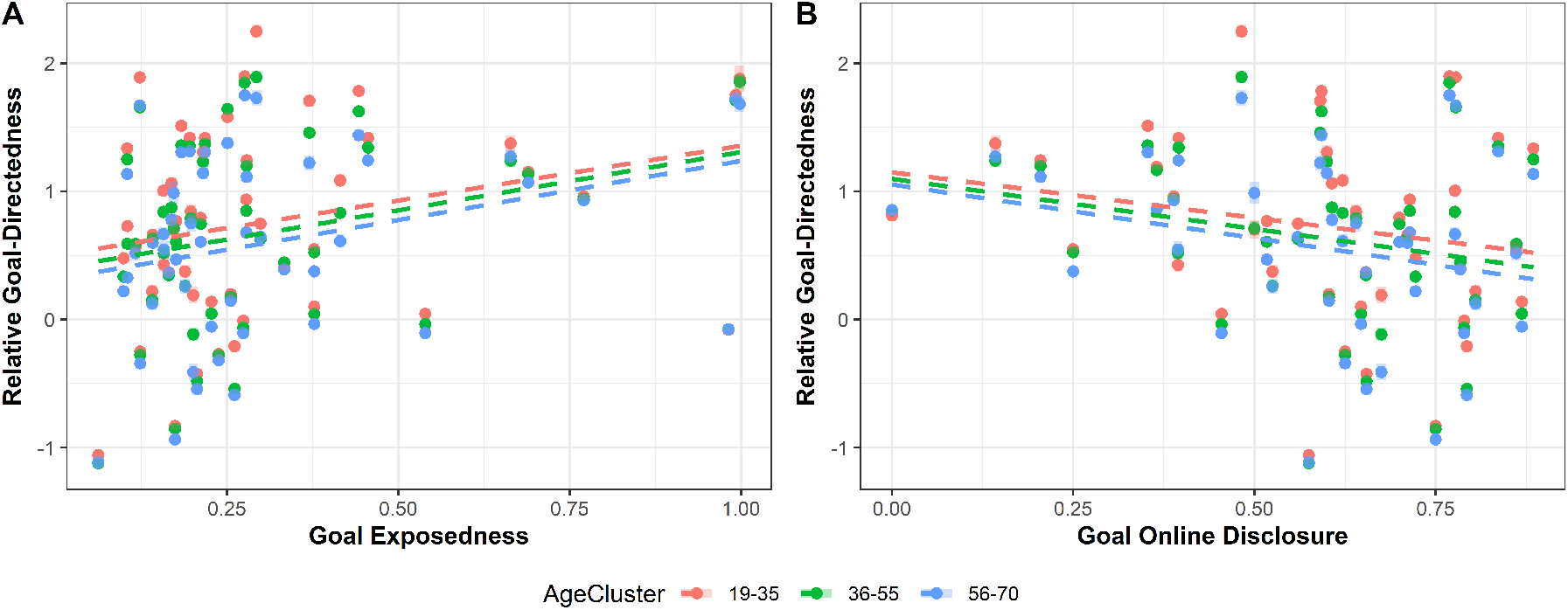
Influence of Goal Exposedness and Goal Online Disclosure on Relative Goal-Directedness Across Age Clusters. A) Scatter plot illustrating the relationship between Relative Goal-Directedness and Goal Exposedness. Each dot represents a level, color-coded by age cluster. The dashed lines represent the linear regression fits for each age cluster, showing a positive trend between Relative Goal-Directedness and Goal Exposedness across all age groups. B) Scatter plot depicting the relationship between Relative Goal-Directedness and Goal Online Disclosure. Each dot represents a level, color-coded by age cluster. The dashed lines show the linear regression fits for each age cluster, indicating a slight decrease in Relative Goal-Directedness with increasing goal online disclosure. The data suggests that age clusters exhibit similar trends in Relative Goal-Directedness relative to goal characteristics.

### A.7 Supplementary tables

**Table S1:** ANOVA Summary Results for each condition shown in Figure 6.

| $N_{goals}$ | Goal ID | $F_{value}$ | DoF | $p_{value}$ |
| --- | --- | --- | --- | --- |
| 3 | 1 | 92097.0684 | 3 | $\ll 10^{-3}$ |
| 3 | 2 | 167567.9079 | 3 | $\ll 10^{-3}$ |
| 3 | 3 | 219396.6102 | 3 | $\ll 10^{-3}$ |
| 4 | 1 | 81737.4851 | 4 | $\ll 10^{-3}$ |
| 4 | 2 | 72731.2548 | 4 | $\ll 10^{-3}$ |
| 4 | 3 | 52497.7072 | 4 | $\ll 10^{-3}$ |
| 4 | 4 | 74606.1757 | 4 | $\ll 10^{-3}$ |

## Notes

### Competing Interest Statement

The authors have declared no competing interest.

### Summary of Updates

Additional analyses to assess the separability of motor and cognitive components in navigation performance

